# Resting-state and task-related neurometabolite levels differentially relate to cortical excitability in the sensorimotor and prefrontal cortex

**DOI:** 10.64898/2026.01.23.701225

**Authors:** Marten Nuyts, Sima Chalavi, Geraldine Rodriguez-Nieto, Koen Cuypers, Raf Meesen, Stephan P. Swinnen, Sybren Van Hoornweder

## Abstract

**Background:** Normal brain function requires a dynamic balance between inhibition and excitation. While magnetic resonance spectroscopy (MRS) quantifies the chief inhibitory and excitatory neurometabolites, GABA and glutamate-glutamine (Glx), the combination of transcranial magnetic stimulation and electroencephalography (TMS-EEG) provides a complementary measure of cortical inhibition-excitation dynamics via transcranial evoked potentials (TEPs). However, the relationship between neurometabolite concentrations and TEPs is unclear.

**Objective:** To characterize the relationship between neurometabolite concentrations and TEPs, as a function of TEP component, TMS pulse type, brain region, neurometabolite, and MRS brain state.

**Methods:** Twenty-five young healthy adults completed a 4-day protocol. Sessions 1 and 2 involved screening, anatomical MRI, and functional MRI localization of DLPFC. In session 3, single- and paired-pulse TMS-EEG were applied over SM1 and DLPFC. Session 4 included resting-state and motor-task-related MRS of SM1 and DLPFC.

**Results:** In SM1, task-related GABAergic tone strongly predicted early to mid-latency TEPs. In DLPFC, local early to mid-latency TEPs showed no relationship to neurometabolites, whereas late and global TEP outcomes revealed some links. The later N100 TEP was the only component consistently modulated by paired-pulse TMS. Task-related MRS measures consistently outperformed resting-state measures in predicting TEPs for SM1, while the opposite was true for DLPFC.

**Conclusions:** Single-pulse TEPs reliably index the local inhibitory tone in SM1, with limited added value from paired-pulse paradigms. These findings support the use of single-pulse TEPs as accessible markers of cortical inhibition, especially for SM1, and may inform biomarkers and strategies for individualized neuromodulation.

**Key point summary:**

- Normal brain function relies on a balance between inhibition and excitation, yet the relationship between local cortical neurometabolite levels and cortical excitability in humans remains poorly understood.
- We measured inhibitory and excitatory neurometabolites using magnetic resonance spectroscopy and assessed cortical excitability responses to non-invasive stimulation using combined transcranial magnetic stimulation - electroencephalography (TMS-EEG).
- In the primary sensorimotor cortex (SM1), local cortical responses to stimulation were best explained by inhibitory GABA levels during motor task performance. In the dorsolateral prefrontal cortex (DLPFC), global cortical responses were best predicted by resting-state neurometabolite levels.
- Overall, paired-pulse TMS paradigms offered little additional value over single-pulse paradigms for informing on neurometabolite levels.
- By demonstrating region- and state-dependent links between neurometabolite levels and cortical excitability, our findings position TMS-EEG as an accessible biomarker of cortical inhibition, particularly in SM1.

*Graphical Abstract:* 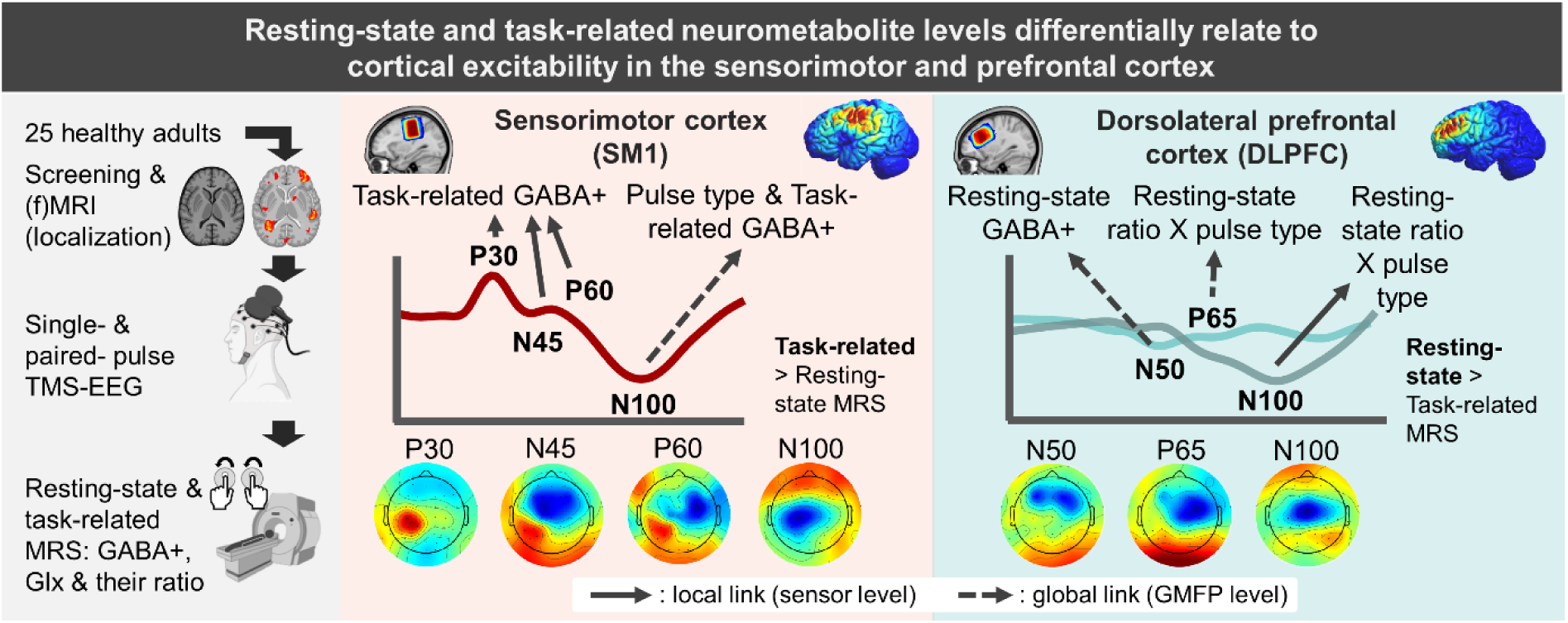

*Abstract figure legend:* The left panel provides a stepwise overview of the study protocol, including (i) participant screening, (ii) functional and anatomical magnetic resonance imaging (MRI) scanning for individualization purposes, (iii) single- and paired-pulse transcranial magnetic stimulation–electroencephalography (TMS-EEG), and (iv) magnetic resonance spectroscopy (MRS) acquired at rest and during a bimanual motor task. The middle and right panels show the grand-average EEG responses to single-pulse TMS over the sensorimotor cortex (SM1, red) and dorsolateral prefrontal cortex (DLPFC, blue), respectively. The topographical maps below each EEG response depict the averaged scalp distributions corresponding to each canonical transcranial evoked potential. Per region, the top left illustration shows the group-level MRS voxel placement, whereas the top right illustration shows the simulated TMS-induced electric fields. While early to mid-latency local cortical responses were best predicted by task-related GABAergic inhibition in SM1, global cortical responses following DLPFC stimulation were best predicted by resting-state neurometabolite levels.

## 1. Introduction

A defining feature of the brain is its ability to maintain a dynamic balance between inhibitory and excitatory influences. Gamma-aminobutyric acid (GABA) mediated inhibition and glutamatergic excitation jointly regulate neuronal firing, shape cortical oscillations and coordinate distributed network activity. When this balance shifts, cognition, perception and motor behavior can be profoundly disrupted, as seen in a wide array of psychiatric and neurological conditions (Sohal & Rubenstein, 2019; Ghatak *et al*., 2021; Li *et al*., 2025). This has fueled interest in developing non-invasive methods to quantify inhibitory and excitatory function, both to understand the healthy brain and to identify biomarkers for diagnosis, prognosis, and personalized treatments.

Magnetic resonance spectroscopy (MRS) and transcranial magnetic stimulation (TMS) are among the most widely used techniques to probe cortical inhibition and excitation. MRS enables localized in-vivo quantification of brain neurometabolites, including GABA, the glutamate-glutamine complex (Glx), and their ratio. Task-related MRS extends this approach by measuring metabolite concentrations during task performance, revealing state-dependent metabolic dynamics linked to specific motor or cognitive demands (Cuypers & Marsman, 2021; Pasanta *et al*., 2023; Li *et al*., 2024).

TMS provides a complementary, neurophysiological, perspective. A single magnetic pulse synchronously excites cortical neurons, leading to a transsynaptic propagation of neural activity along the brain (Siebner *et al*., 2022). Traditional single-pulse TMS over the primary sensorimotor cortex (SM1) yields indirect indices of corticospinal excitability, via motor evoked potentials, as assessed by means of electromyography (Spampinato *et al*., 2023; Nuyts *et al*., 2025b). Paired-pulse TMS paradigms probe GABAergic and glutamatergic processes more selectively, with long-interval intracortical inhibition (LICI) TMS indexing the slower, longer-lasting, GABA_B_ receptor system, as compared to the faster, more transient, GABA_A_ system (Valls-Solé *et al*., 1992; Werhahn *et al*., 1999). However, electromyography-based TMS work captures only the final corticospinal output, and is therefore limited to SM1, minimizing its ability to isolate cortical mechanisms and generalize to non-motor regions such as the dorsolateral prefrontal cortex (DLPFC), a key region implicated in neuropsychiatric disorders and brain stimulation interventions.

These limitations are largely overcome by combining TMS with electroencephalography (TMS-EEG), allowing to more directly investigate cortical GABAergic and glutamatergic processes (Ilmoniemi & Kicić, 2010). Following a single TMS-pulse, a series of transcranial evoked potentials (TEPs) captured by EEG reflect the spatiotemporal signal propagation across local and remote brain regions (Ilmoniemi *et al*., 1997; Hernandez-Pavon *et al*., 2023). Pharmacological work over SM1 suggests that earlier (< 50 ms) components are mediated by GABA_A_, whereas later components reflect GABA_B_ activity (Premoli *et al*., 2014a; Darmani & Ziemann, 2019). Paired-pulse TMS-EEG further enables the quantification of cortical inhibitory dynamics linked to GABA_B_ receptors in both SM1 (Farzan *et al*., 2010; Premoli *et al*., 2014b; de Goede *et al*., 2020) and DLPFC (Daskalakis *et al*., 2008; Farzan *et al*., 2010; Salavati *et al*., 2018; Poorganji *et al*., 2023; Takano *et al*., 2025).

Despite conceptual links between MRS-derived neurometabolites and TMS-derived cortical excitability, empirical evidence connecting both modalities remains scarce and inconsistent. Most research has reported null associations between TMS-derived GABAergic synaptic activity and MRS GABA concentrations (for a review, see (Cuypers & Marsman, 2021)). Conversely, several studies have demonstrated links between TMS measures and Glx (Stagg *et al*., 2011; Tremblay *et al*., 2013; Dyke *et al*., 2017).

However, a major limitation of much prior work is its reliance on electromyography as a readout of TMS effects, which conflates cortical and spinal contributions and restricts inference largely to the motor cortex (Spampinato *et al*., 2023). To our knowledge, TMS-EEG and MRS have been integrated only once, where the late N100 TEP component was shown to correlate to both GABA and Glx levels in the left prefrontal cortex, but not in the motor cortex (Du *et al*., 2018). However, late TEP components such as the N100 are highly prone to multisensory co-activation, complicating their neurophysiological interpretation (Conde *et al*., 2019).

Here, we investigated neurometabolite-cortical excitability relationships using combined MRS and TMS-EEG. We aimed to identify cortical measures that may serve as sensitive, non-invasive biomarkers of inhibitory and excitatory processes. We acquired GABA and Glx concentrations from left SM1 and DLPFC during rest and a motor coordination task via MRS. Cortical excitability was indexed as TEPs, generated by single-pulse and paired-pulse (LICI) TMS-EEG, with the latter providing a readout of GABA_B_ activity.

We hypothesized that neurometabolite-cortical excitability relationships are context-dependent, varying as a function of TEP component, TMS pulse type (single vs. paired-pulse TMS), brain region (SM1 vs. DLPFC), neurometabolite (GABA vs. Glx), and MRS brain state (rest vs. task). Specifically, we expected stronger neurometabolite associations for early to mid-latency TEP components than for the later N100, given its susceptibility to multisensory co-activation (Conde et al., 2019). We did predict a selective modulation of N100 by the paired-pulse LICI paradigm with respect to single-pulse TMS in a GABA-dependent manner due to its shared links to GABA_B_ activity (Premoli *et al*., 2014b). We further expected robust neurometabolite-TEP associations in SM1, in line with pharmacological TMS-EEG findings (Darmani & Ziemann, 2019), whereas no strong hypotheses were formulated for DLPFC due to its more limited pharmacological characterization (Rogasch *et al*., 2020). Concerning neurometabolite specificity, prior TMS-MRS studies have more consistently reported Glx associations, but these findings were largely based on corticospinal output (Cuypers & Marsman, 2021). We therefore considered it plausible that GABA, or the balance between GABA and Glx, better predict TEPs, in line with pharmacological work, and accordingly did not formulate any hypotheses (Darmani & Ziemann, 2019; Belardinelli et al., 2021). Finally, we expected stronger coupling between task-based MRS measures and TMS-EEG outcomes than during resting-state MRS, particularly in SM1, given the motor nature of the task.

## 2. Methods

### 2.1. Participants

Twenty-five healthy volunteers (12 females; mean age: 22; range: 18-31 years) were enrolled. Participants had (corrected-to-)normal vision, and were right-handed as assessed by the Oldfield Handedness Inventory (Oldfield, 1971) (mean lateralization quotient 89.21 ± SD: 12.87). Also, they had no contraindications for TMS or magnetic resonance imaging (MRI) and reported no history of neurological diseases or psychiatric disorders. The Montreal Cognitive Assessment (MoCA; (Nasreddine *et al*., 2005); cut-off score ≤ 26) was used to assess cognitive and executive functioning (mean score: 28.52 ± SD: 0.92).

All participants provided written informed consent in accordance with the Declaration of Helsinki. The protocol was approved by the regional ethical committee (KU Leuven, Belgium; protocol number: S63320).

### 2.2. Experimental procedure

The current work was part of a larger six-session research protocol that stretched over a mean time of 18.53 (± SD: 5.87) days. Here, we focus on four sessions (for an overview, see **Figure 1**).

**Figure 1.**
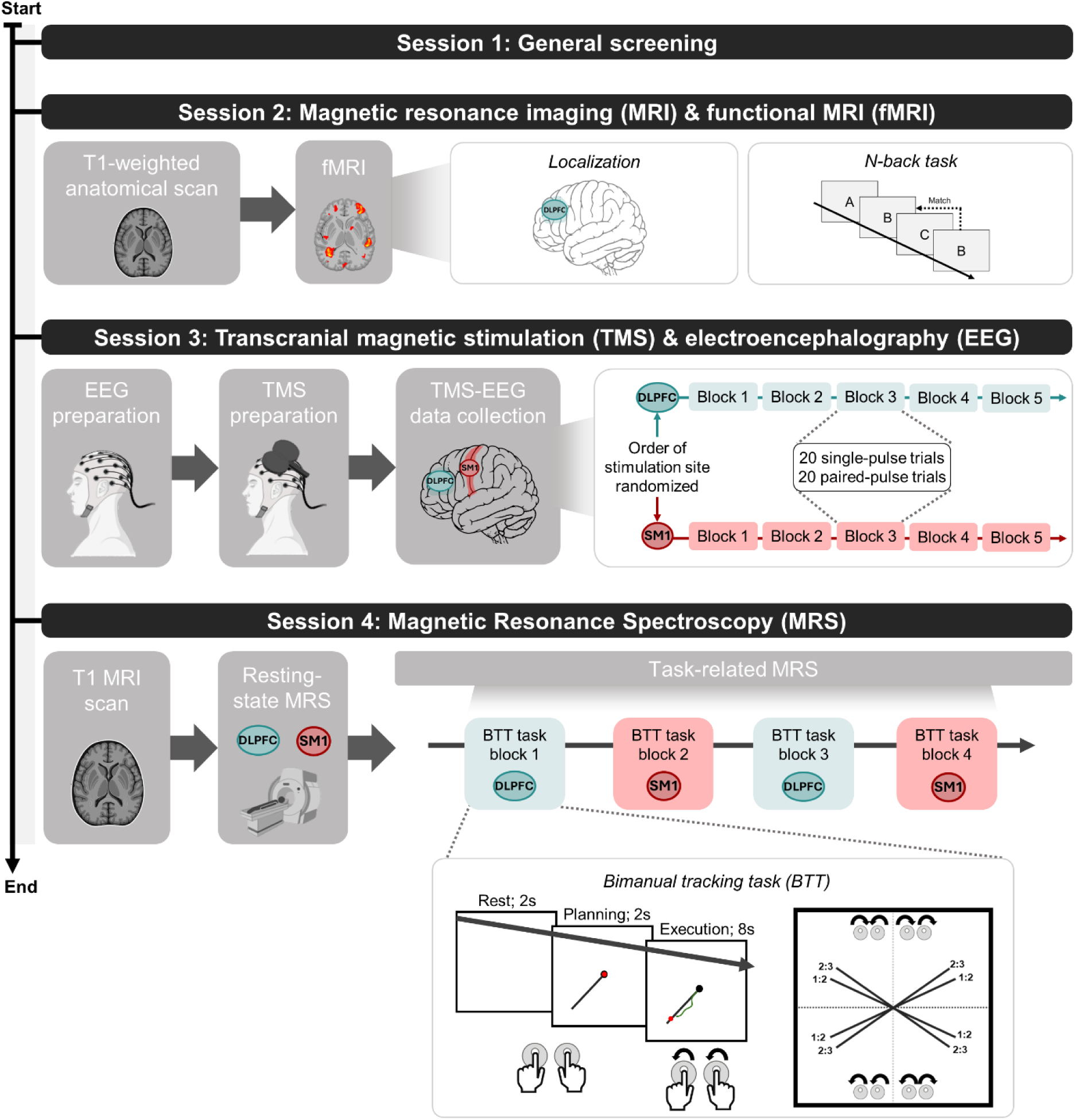
A top-down chronological overview of the study protocol and its measurement tools. The study consists of four sessions. Session 1 consisted of eligibility screening, followed by structural and functional magnetic resonance imaging (MRI) scanning in session 2. Session 3 consisted of transcranial magnetic stimulation-electroencephalography (TMS-EEG) over the left primary sensorimotor cortex (SM1) and the left dorsolateral prefrontal cortex (DLPFC). Finally, session 4 consisted of resting-state magnetic resonance spectroscopy (MRS) and task-related MRS during performance of the bimanual tracking task (BTT).

In session 1 (day 1), participants completed several questionnaires regarding potential contra-indications, handedness, and cognitive and executive functioning.

In session 2 (day 2), a structural MRI scan was acquired, which was later uploaded into the TMS neuronavigation software. Also, participants performed a working memory task while functional MRI (fMRI) data were collected. The obtained functional data served to localize left DLPFC during TMS-EEG and MRS data collection sessions.

In session 3 (day 3), TMS-EEG data were collected over left SM1 and DLPFC using an intermixed single- and paired-pulse TMS protocol. The stimulation order was randomized across participants.

Lastly, in session 4 (day 4 or 5), MRS data were collected over left SM1 and DLPFC at rest and while performing a bimanual motor coordination task, the bimanual tracking task (BTT) (Sisti *et al*., 2011). For thirteen participants, session 4 occurred on day 4, and for the remaining twelve, on day 5. This discrepancy was due to the broader research protocol studying two distinct versions of the bimanual motor coordination task (see further), whereas the current work solely focusses on the version which contains the largest number of participants. **Supplementary materials 1** shows that the MRS measurements over the two sessions correlate well (all p-values < 0.02).

Ultimately, TMS-EEG data were not collected for one participant due to technical issues, and several participants were excluded based on (TMS-EEG) data quality. This resulted in a complete dataset of 20 and 14 participants for SM1 and DLPFC, respectively.

### 2.3. Functional magnetic resonance imaging (fMRI)

All (f)MRI and MRS data were acquired on a 3T Philips Achieva scanner (University hospital Leuven, Gasthuisberg), with a 32-channel receiver head coil (Philips, Best, The Netherlands). On Day 2, a T1-weighted high-resolution scan was collected using a three-dimension turbo field echo (3DTFE) (TR = 9.7 ms, TE = 4.6 ms, flip angle = 8°, voxel size = 1 x 1 x 1 mm3, field of view (FoV) = 256 × 242 × 182 mm^3^, 182 sagittal slices, scan time = ∼ 6 min). The individual’s T1-weighted image was later uploaded to the TMS neuronavigation software. Next, fMRI data were collected with an ascending gradient EPI pulse sequence for T2*-weighted images (TR = 2000 ms, TE = 29.80 ms, multiband factor 2, flip angle = 90°, voxel size = 2.5 × 2.5 × 2.5 mm^3^, FoV = 210 × 210 × 145.6 mm^3^, 54 transverse slices, interslice gap = 0.2 mm, 171 dynamical scans, scan time ∼ 6 min) while participants performed an N-back task (see **Supplementary Materials 1** for task details).

### 2.4. Electroencephalography (EEG) and electromyographic recordings

EEG data were recorded using BrainVision Recorder software (Brain Products GmbH, Germany) from 64 actiCAP slim active electrodes attached to an EEG cap (10-20 system, Brain Products GmbH, Germany) with the ground and reference electrodes placed on the forehead (electrode Fz) and the posterior center of the head (electrode CPz), respectively. A TMS-compatible EEG amplifier (actiCHamp Plus 64 System, Brain Products GmbH, Germany) was used with a sampling frequency of 10 kHz. Impedance levels were continuously monitored and kept below 20 kΩ. Participants were seated comfortably and instructed to remain relaxed, minimize their blinking, and keep their eyes open.

Electromyographic data were recorded from the right first dorsal interosseous muscle using self-adhesive surface Ag-electrodes (Bagnoli-16, Delsys Inc, Boston, USA) placed on the prepared skin. The reference electrode (enraf-en-trode, Enraf-Nonius, Amsterdam, NL) was placed on the right distal ulnar bone. A HumBug Line Noise Eliminator (Quest Scientific, North Vancouver, BC, Canada) was used, and the data were amplified (gain = 1000), bandpass filtered (4-1500 Hz), and eventually digitized at 5 kHz (CED Signal Version 4.11, Cambridge Electronic Design, Cambridge, UK).

### 2.5. Transcranial Magnetic Stimulation (TMS)

Per brain region, monophasic TMS pulses were delivered every 5 s (+/- 10% jitter) using a 70 mm figure-of-eight coil coated with a thin foam layer, connected to two Magstim 200 units (Magstim Company, Dyfed, UK). The coil was placed perpendicular to the head and oriented 45° relative to the midline, with the coil handle pointing posterolateral, inducing a posterior-anterior current in the brain. The TMS coil’s position was continuously monitored throughout the experiment via stereotaxic navigation (Visor 2, ANT Neuro) using the T1-weighted MRI scan.

For left DLPFC, the coil was placed over the mask of peak fMRI activity during the N-back task, which was uploaded to the neuronavigation software. For left SM1, we searched for the optimal stimulation site by moving the coil in ∼1 cm steps to locate the motor hotspot, defined as the location where the largest and most consistent motor evoked potential amplitudes in the right first dorsal interosseous muscle were obtained. Over the motor hotspot, we determined the resting motor threshold, defined as the minimal intensity required to elicit a motor evoked potential with a minimal peak-to-peak amplitude of 50 µV in at least five out of ten trials.

The stimulation intensity for both stimulation sites was set to the intensity which evoked an average motor evoked potential amplitude of 1 mV in five consecutive TMS pulses over SM1. Per brain region, a total of 100 single and paired TMS pulses were delivered intermixed, split up across five recording blocks (lasting ∼200 s each). Paired-pulse trials consisted of a LICI protocol using conditioning and test stimuli set to an intensity of 1 mV, with an interstimulus interval of 100 ms. During data collection, participants wore earphones (with earmuffs) that administered white noise to minimize the effect of the TMS clicking sound.

### 2.6. Magnetic Resonance Spectroscopy (MRS)

Prior to MRS scanning, a short T1-weighted scan was collected for MRS voxel placement (TR = 9.6, TE = 4.6 ms, flip angle = 8°, voxel size = 1.5 x 1.5 x 1.5 mm^3^, FoV = 256 x 256 x 182 mm^3^, 182 sagittal slices, scan time = ∼ 2 min). For MRS, a GABA-edited MEGAPRESS sequence was used to quantify GABA and GLx levels (TR = 2000 ms, TE = 68 ms, flip angle = 90°, 2 kHz spectral width, ‘ON’/’OFF’ editing pulse frequency = 1.9 / 7.5 ppm, ‘ON’/’OFF’ spectra collected in an interleaved fashion) (Mescher *et al*., 1998; Edden & Barker, 2007). Automatic shimming was performed before every MRS acquisition, and 16 unsuppressed water averages were obtained over each brain region of interest and later used for referencing.

Acquisition voxels were placed on an individual basis over left DLPFC (voxel size = 40 × 25 × 25 mm^3^, 160 averages, scan time = ∼ 11 min) and left SM1 (voxel size = 30 × 30 × 30 mm^3^, 112 averages, scan time = ∼ 8 min). Specifically, the DLPFC voxel was placed over the mask of peak fMRI activity during the N-back task, while the SM1 voxel was placed over the left hand knob area of the precentral gyrus (Yousry *et al*., 1997). Of note, due to the contribution of macromolecules at 3 ppm, we refer to GABA+ in all future analyses.

MRS data were acquired both at rest (i.e., resting-state MRS) and during a motor coordination task, the bimanual tracking task (BTT) (i.e., task-related MRS). Resting-state MRS was collected before task-related MRS data. During resting-state MRS, participants were instructed to relax and keep their eyes fixed at a cross.

Task-based MRS was collected in four blocks (two per brain region) while participants performed the BTT (see **Supplementary Materials 1** for task details). In this task, participants tracked a moving cue on the screen as accurately as possible. The trial consisted of three stages (cf., **Figure 1**), rest, planning and execution, and engaged both prefrontal and sensorimotor areas (Swinnen & Wenderoth, 2004; Beets *et al*., 2015; Santos Monteiro *et al*., 2017). During the planning stage, participants observed the cue but were not allowed to move. During execution, they performed continuous bimanual rotations with the index fingers, with direction and speed varying across conditions. The BTT included 8 distinct conditions that manipulated inter-hand frequency (relative speed of both hands, either a 1:2 or 2:3 ratio) and movement direction, with both indices either rotating left, right, inwards, or outwards.

To maintain accurate voxel placement throughout the session, additional short T1-weighted scans were acquired prior to the first and third task block. These scans allowed verification and adjustment of the voxel in case of head movement.

### 2.7. Data analysis

EEG data were preprocessed and analyzed using custom MATLAB scripts (2022b, The Math-Works Inc., Portola Valley, California, USA) using EEGLAB (Delorme & Makeig, 2004) and Fieldtrip functions (Oostenveld *et al*., 2011).

Data were epoched from -500 to 500 ms around the TMS pulse, which was the test pulse for paired-pulse TMS. Next, the TMS pulse artifact was removed and interpolated (cubic) from -5 ms to 20 ms (SM1) or 30 ms (DLPFC). Then, the data was filtered (2^nd^ order zero-phase Butterworth filter at 1-49 Hz), downsampled to 500 Hz, re-referenced to the common average, baseline corrected from -110 ms to -10 ms for single-pulses and -210 to -110 for paired-pulses, and finally averaged. Single trials were visually checked for blinks or muscle activity and removed accordingly (median: 17). Channels containing noise or suffering from extensive decay artifacts were interpolated (median: 9).

Of note, in paired-pulse TMS-EEG paradigms, the response to the test stimulus is intermixed with activity evoked by the preceding conditioning stimulus (Daskalakis *et al*., 2008). To isolate the test-pulse–specific response and facilitate comparison with single-pulse TEPs, the averaged single-pulse TEP was subtracted from the averaged paired-pulse response time-locked to the conditioning stimulus (for details, see (Premoli *et al*., 2014b)).

Prototypical TEP peaks, defined based on their latency (in ms) and polarity (positive or negative), were extracted at the single-channel level, with the channel selected according to the group-averaged topographical distribution for each TEP component. For SM1, electrode C3 was used for all TEP peaks (P30, N45, P60, and N100). For DLPFC, electrode F3 was used for the mid-latency TEP peaks (N50 and P65) and electrode Cz for the latest TEP component (N100).

In addition to these local, electrode-specific measures, we computed the global mean field potential (GMFP). GMFP reflects the spatial standard deviation of EEG amplitudes across all electrodes, providing a summary measure of TEPs independent of electrode locations. GMFP was included as a complementary, location-free robustness check (cf., 2.8. Statistical analyses) (Lehmann & Skrandies, 1980).

MRS data processing was done using the GABA+ analysis Toolkit Gannet (version 3.2.1) (Edden & Barker, 2007). The filled-in minimum reporting standards for in-vivo MRS checklist can be found in **Supplementary Materials 2** (Lin *et al*., 2021). Frequency and phase offsets were corrected using spectral registration (Mikkelsen *et al*., 2020). Next, the difference spectrum of “ON” from “OFF” spectra was fitted using a three-Gaussian function between 2.8 and 4.2 ppm. Water signals were modeled using a Lorentz-Gaussian lineshape and served as the reference metabolite for remaining concentration estimates. MRS voxels were partial-volume corrected by coregistering them to the T1-weighted scan using the GannetCoRegister module (SPM12-VBM8 toolbox) and by identifying fractions of grey matter, white matter, and cerebrospinal fluid within the voxel using the GannetSegment module. Of note, the high-resolution T1-weighted scan acquired on Day 2 was co-registered to the (short) anatomical scans that were used to verify MRS voxel placement, ensuring a more accurate segmentation. Finally, neurometabolite levels were quantified and corrected for different tissue fraction weighting (see Equation 5 (Harris *et al*., 2015)) using the GannetQuantify module. All MRS spectra were visually assessed for poor water suppression (Multiply Optimized Insensitive Suppression Train; MOIST), lipid contamination and by examining the fit error (cut-off: fit error > 10) and signal-to-noise ratio to ensure sufficient data quality.

### 2.8. Statistical analysis

The current work set out to unveil the relationship between several canonical TEP components and neurometabolites. Therefore, we applied single- and paired-pulse TMS-EEG over two regions and obtained GABA+ and Glx data from both regions during two brain states: resting-state and task-performance. We also investigated the GABA+/Glx ratio as an exploratory index to the inhibitory/excitatory balance.

To probe the link between MRS-neurometabolites and TMS-derived cortical excitability for SM1, we examined if TEP components were best predicted by neurometabolites at rest, during execution of a motor task, or by the neurometabolite ratio during rest or task execution. We established four linear mixed effect models per TEP Component (EEG sensor C3: P30, N45, P60, and N100);

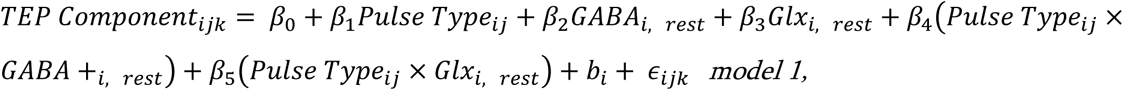

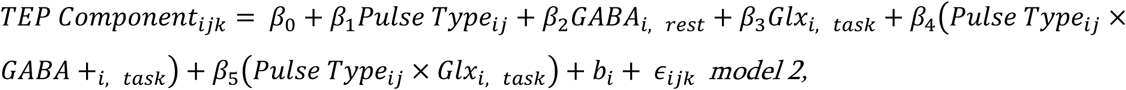

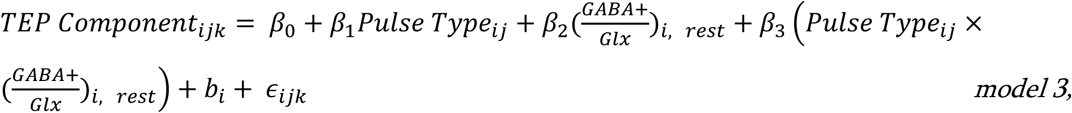

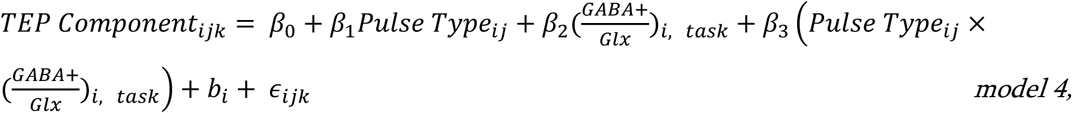

where TEP component_ijk_ denotes the amplitude per TEP component k, for participant i and pulse type j. Participant-specific variability was modeled via a random intercept b_i_.

Following backwards model building, these models were compared in terms of Akaike Information Criterion (AIC), Bayesian Information Criterion (BIC) and R^2^ values.

To probe the link between TEPs and neurometabolites in the left DLPFC, we repeated these analyses for each TEP component following DLPFC stimulation (EEG sensor F3: N50 and P65; EEG sensor Cz: N100).

To complement the local sensor-level peak analyses, we repeated the models using GMFP data as a location-free robustness check. These results are reported in **Supplementary Materials 3** and briefly mentioned in the Results section.

## 3. Results

### 3.1. Task-related GABAergic tone in SM1 predicts early and mid-latency TEP components

**Figure 2** shows the descriptive data related to SM1, including both the neurometabolite and TEP data, as well as a current flow model visualizing the induced electric field following SM1 stimulation.

**Figure 2.**
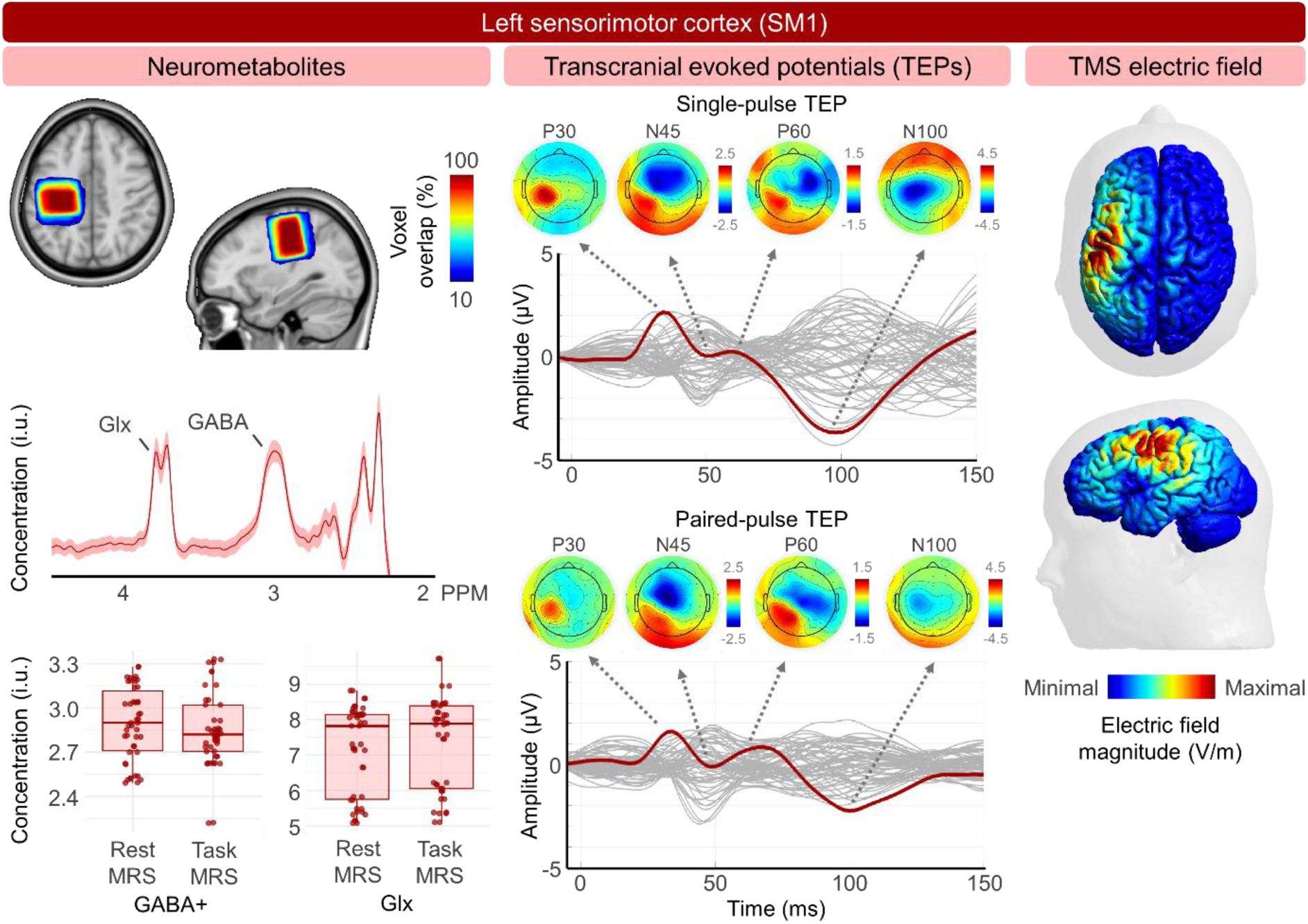
Descriptive data over the left primary sensorimotor cortex (SM1). **Left panel:** Mean group-level voxel overlap (% of total sample size) for magnetic resonance spectroscopy (MRS), mean MR spectra for one measurement timepoint, and descriptive boxplots containing the concentration of GABA+ and Glx during resting-state and task-related MRS. During task-related MRS, participants performed a bimanual motor task. **Middle panel**: Butterfly plots showing the group averaged transcranial evoked potentials (TEPs) induced by a single (top) and paired-pulse (bottom) TMS protocol over left SM1, as captured by electroencephalography (EEG). Per plot, the mean response of all EEG electrodes is shown in grey, with the electrode used for our sensor-level analyses, sensor C3, highlighted in dark red. The TEP traces reflect a series of positive (P) and negative (N) components at different latencies. Four TEP components were identified, P30, N45, P60 and N100. For each of these components, the matching topographies are plotted above. **Right panel:** Visualization of the electric field magnitude induced by transcranial magnetic stimulation (TMS) over the left SM1, made with SimNIBS in a template head model (Thielscher *et al*., 2015).

Across TEP components and models for the sensor level data, pulse type and its interactions were consistently non-significant, indicating that variability in SM1 TEPs was not driven by the type of TMS pulse paradigm. The results of the models are shown in **Table 1** and **Figure 3**. To verify the robustness of these findings, we repeated all analyses using the GMFP signal (**Supplementary Materials 3**).

**Figure 3.**
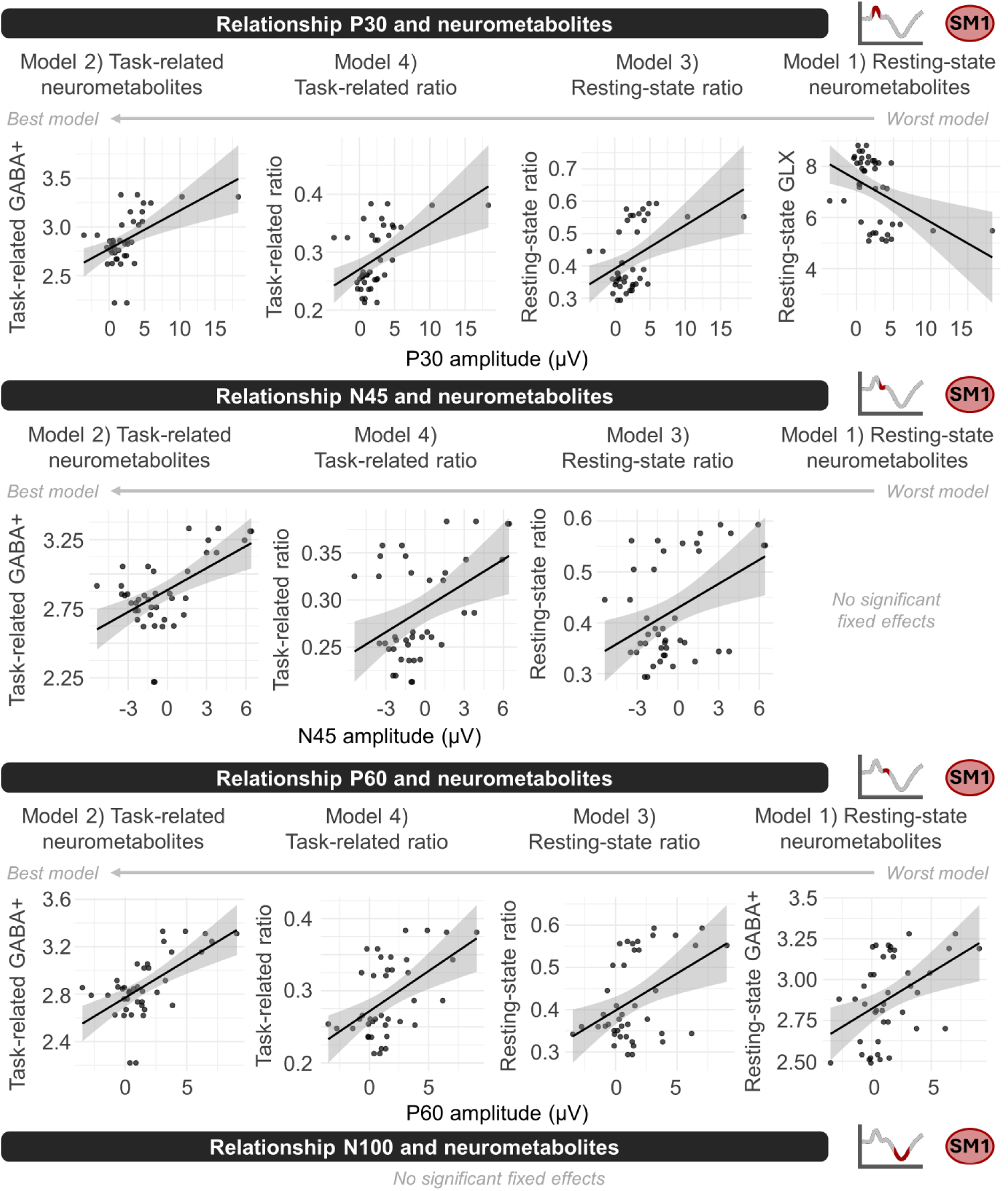
The link between neurometabolites and transcranial magnetic stimulation (TMS) per transcranial evoked potential (TEP) component, extracted from electrode C3 (∼ primary sensorimotor cortex [SM1]). For P30, N45 and P60, multiple models yielded significant fixed effects. For each of these components, the models were ordered from left to right, with left being the best model (cf., Table 1). Model 2, including task-related neurometabolites, and in particular GABA+ during motor task performance, consistently outperformed the other models for P30, N45 and P60. Likewise, task-related MRS measures consistently outperformed resting-state MRS measures. For N100, no significant effects remained across all four models following model building. The GMFP analyses (**Supplementary Materials 3**) mostly concur with the sensor level data, with the largest difference being a significant effect of pulse type on N100.

**Table 1.**
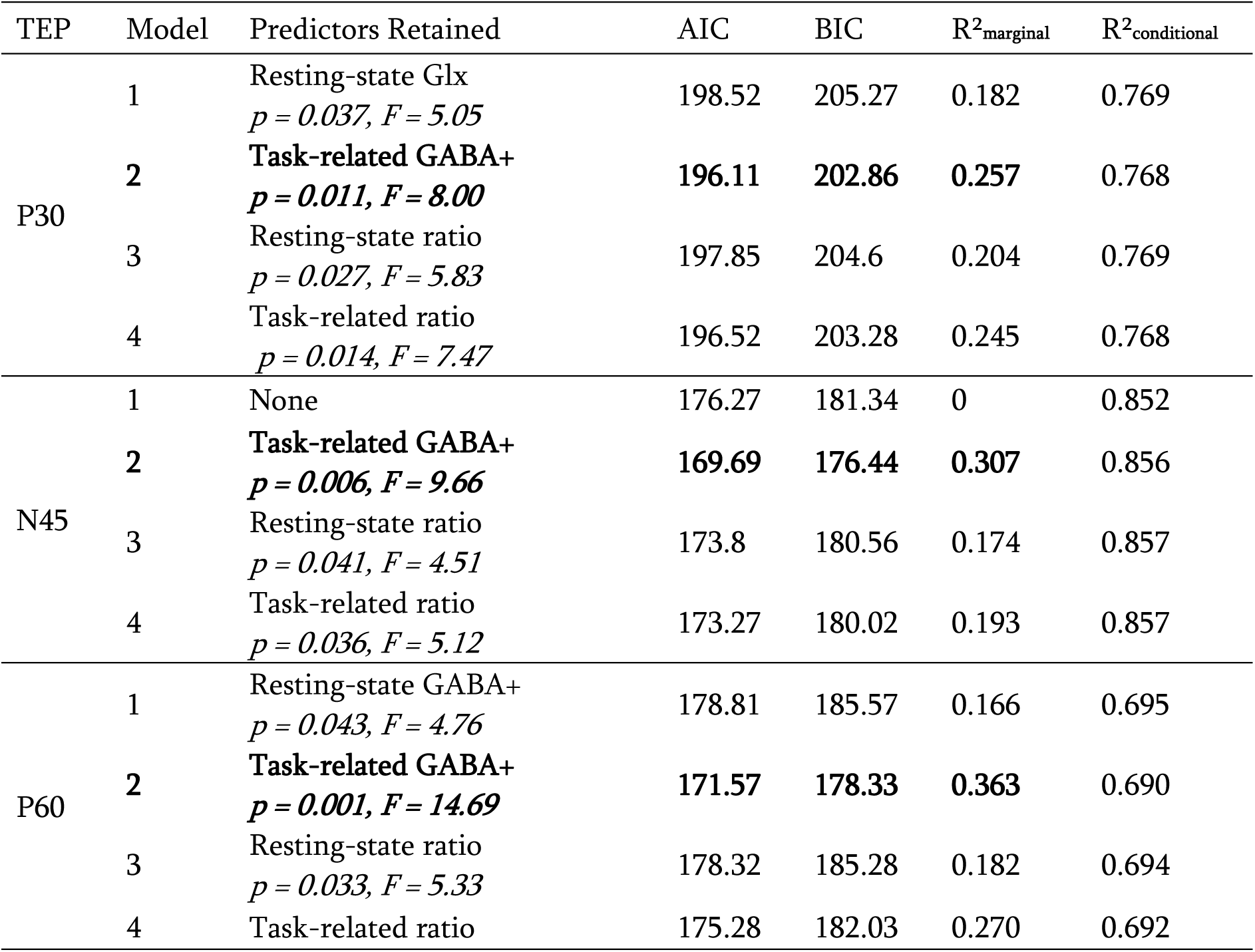
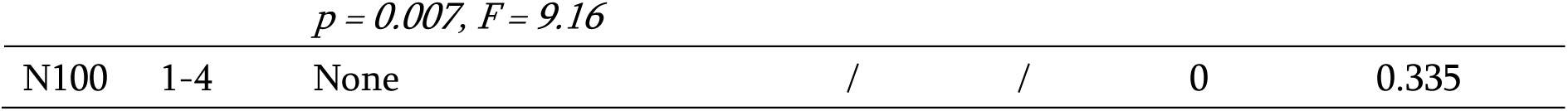
Results of models examining the link between SM1 TEP components and neurometabolites.

| TEP | Model | Predictors Retained | AIC | BIC | R<sup>2</sup><sub>marginal</sub> | R<sup>2</sup><sub>conditional</sub> |
| --- | --- | --- | --- | --- | --- | --- |
| P30 | 1 | Resting-state Glx<br>$p = 0.037, F = 5.05$ | 198.52 | 205.27 | 0.182 | 0.769 |
| | 2 | **Task-related GABA+**<br>$p = 0.011, F = 8.00$ | **196.11** | **202.86** | **0.257** | 0.768 |
| | 3 | Resting-state ratio<br>$p = 0.027, F = 5.83$ | 197.85 | 204.6 | 0.204 | 0.769 |
| | 4 | Task-related ratio<br>$p = 0.014, F = 7.47$ | 196.52 | 203.28 | 0.245 | 0.768 |
| N45 | 1 | None | 176.27 | 181.34 | 0 | 0.852 |
| | 2 | **Task-related GABA+**<br>$p = 0.006, F = 9.66$ | **169.69** | **176.44** | **0.307** | 0.856 |
| | 3 | Resting-state ratio<br>$p = 0.041, F = 4.51$ | 173.8 | 180.56 | 0.174 | 0.857 |
| | 4 | Task-related ratio<br>$p = 0.036, F = 5.12$ | 173.27 | 180.02 | 0.193 | 0.857 |
| P60 | 1 | Resting-state GABA+<br>$p = 0.043, F = 4.76$ | 178.81 | 185.57 | 0.166 | 0.695 |
| | 2 | **Task-related GABA+**<br>$p = 0.001, F = 14.69$ | **171.57** | **178.33** | **0.363** | 0.690 |
| | 3 | Resting-state ratio<br>$p = 0.033, F = 5.33$ | 178.32 | 185.28 | 0.182 | 0.694 |
|  | 4 | Task-related ratio | 175.28 | 182.03 | 0.270 | 0.692 |
| $$p = 0.007, F = 9.16$$ |  |  |  |  |  |  |
| N100 | 1-4 | None | / | / | 0 | 0.335 |

#### 3.1.1. The P30 component

For P30, across all models, conditional R² values were high (**Table 1**). The best-performing model was model 2, in which task-related GABA+ positively predicted P30 amplitude (**Figure 3**). Models containing the ratio between GABA+ and Glx also explained part of the variance, whereas resting-state neurotransmitters alone were least predictive. GMFP analyses confirmed these findings, with Model 2 again providing the best fit. Notably, the GMFP data revealed a task-related GABA+ × pulse type interaction: higher task-related GABA+ predicted larger P30 amplitudes in the single-pulse condition, but this effect was reduced in the paired-pulse condition, consistent with stronger local inhibitory modulation under high GABA+ levels.

Overall, these results indicate that early P30 amplitudes are strongly predicted by the task-related GABAergic tone, with little modulation by the type of stimulation paradigm. In other words, higher levels of inhibitory neurometabolites seem to be associated with higher TEP P30 peak amplitudes.

#### 3.1.2. The N45 component

All N45 models showed high conditional R² values. Again, the model with task-related GABA+ provided the strongest explanatory power, with larger GABA+ levels during task performance predicting smaller (less negative) N45 amplitudes. Models including GABA+/Glx ratios (task-related and resting-state) also showed significant but weaker associations. Resting-state neurotransmitters did not predict N45 amplitude. GMFP analyses converged with these sensor-based analyses, as N45 was best predicted by task-related GABA+ with task-related GABA+/Glx ratios providing weaker but significant effects.

Together with P30, these findings show that early TEP components are most strongly associated with inhibitory neurometabolites, especially during task engagement.

#### 3.1.3. The P60 component

P60 showed a highly similar pattern. The task-related GABA+ model provided the best fit and explained the largest proportion of variance, indicating that stronger task-related GABAergic tone was associated with larger P60 amplitudes. Task-related and resting-state GABA+/Glx ratios were also predictive, whereas resting-state neurometabolite concentrations explained the least variance, with a significant effect of resting-state GABA+ concentration. Akin to N45, GMFP analyses converged with these local sensor analyses.

Overall, the P60 results reinforce the interpretation that early to mid-latency TEP components are jointly modulated by GABAergic processes, with task-related inhibitory tone revealing the strongest link. In other words, higher early to mid-latency TEP peak amplitudes were associated with an increased GABAergic inhibitory tone in SM1.

#### 3.1.4. The N100 component

In contrast to the earlier components, none of the models explained variance in N100 amplitude, and no MRS nor TMS pulse type related effects were significant. Task-related GABA+ was, however, close to significance, with a p-value of 0.051, just like pulse type (p = 0.064). Whereas sensor-based results yielded no significant effects, GMFP N100 showed a significant effect of task-related GABA+ (p = 0.009), alongside a strong main effect of pulse type (p < 0.001), with larger (more negative) N100 amplitudes in the single-pulse condition. The dissociation at the sensor level with respect to the other TEP components indicates that N100 reflects distinct processes that may not be captured by SM1 GABA+/Glx measures in either rest or task conditions, albeit that some fixed effects neared the significance threshold. Furthermore, at the global field level, N100 was strongly shaped by the stimulation paradigm in addition to neurometabolites levels. This concurs with the notion that the late N100 response is a broader network and multisensory response rather than solely a region-specific inhibitory tone.

### 3.2. Local DLPFC TEPs are largely independent of local neurometabolites

Consistent with SM1, **Figure 4** shows the descriptive neurometabolite and TEP data for DLPFC, as well as a current flow model visualizing the induced electric field following DLPFC stimulation. Analogous models were applied to DLPFC TEPs (N50, P65, N100). The results of the models are shown in **Table 2** and **Figure 5**.

**Figure 4.**
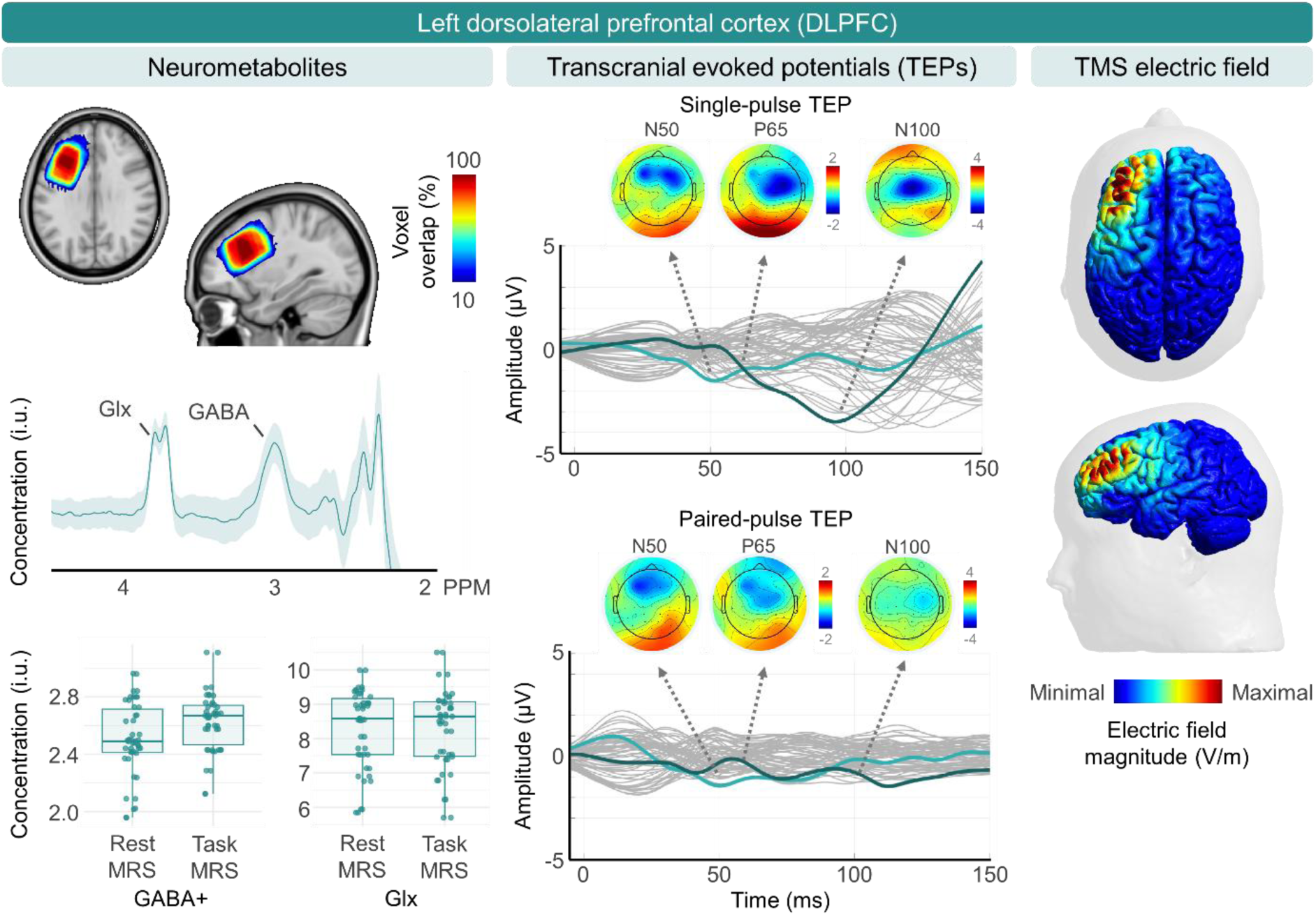
Descriptive data over the left dorsolateral prefrontal cortex (DLPFC). **Left panel:** Mean group-level voxel overlap (% of total sample size) for magnetic resonance spectroscopy (MRS), mean MR spectra, and descriptive boxplots containing the concentration of GABA+ and Glx during resting-state MRS and task-related MRS. During task-related MRS, participants performed a bimanual motor task. **Middle panel**: Butterfly plots showing the group averaged transcranial evoked potentials (TEPs) induced by a single (top) and paired-pulse (bottom) TMS protocol over left DLFPC, as captured by electroencephalography (EEG). Per plot, the mean response of all EEG electrodes is shown in grey, with the electrodes used for our sensor-level analyses, sensors F3 (N50 & P65) and Cz (N100), highlighted in light blue and dark blue, respectively. The TEP traces reflect a series of positive (P) and negative (N) components at different latencies. Three TEP components were identified in the butterfly plots (i.e., N50, P65 and N100). For each of these components, the matching topographies are plotted above. **Right panel:** Visualization of the electric field magnitude induced by transcranial magnetic stimulation (TMS) over the left DLPFC, made with SimNIBS in a template head model (Thielscher *et al*., 2015).

**Figure 5.**
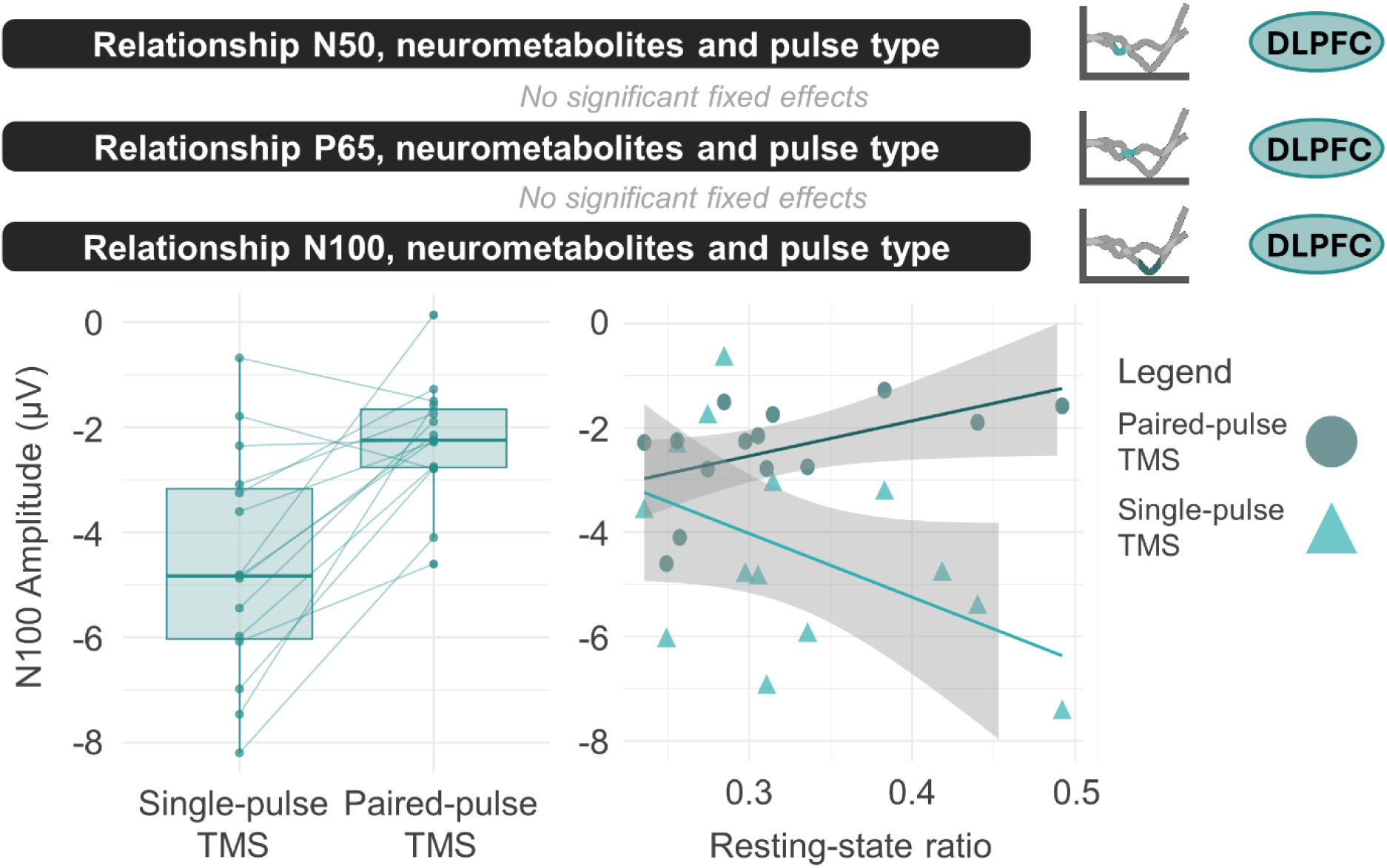
The link between left dorsolateral prefrontal cortex (DLPFC) neurometabolites and transcranial magnetic stimulation (TMS) per transcranial evoked potential (TEP) component, extracted from EEG electrodes F3 (N50 and P65) and Cz (N100). For N50 and P65, no significant effects occurred on the sensor-level data. For N100 (Table 2), an effect of pulse type was observed, with a larger N100 during single-pulse stimulation. Also, an interaction between pulse type and the resting-state ratio between GABA+ and Glx was found. A larger amount of GABA+ relative to Glx was related to a larger N100 only for single-pulse TMS. In the paired-pulse condition, participants with a stronger inhibitory tone had an inhibited response, quantified as a lower N100.

**Table 2.**
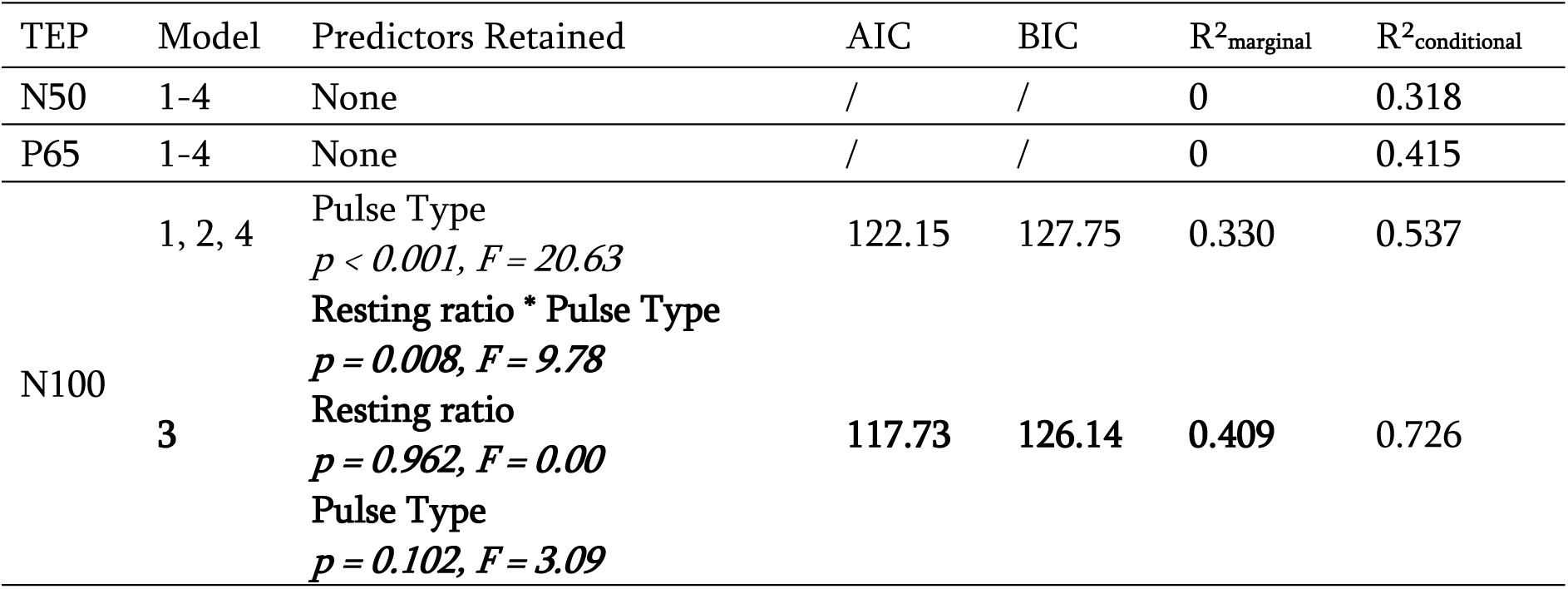
Results of models examining the link between DLPFC TEP components and neurometabolites.

#### 3.2.1. The N50 and P65 components

For the early to mid-latency components, N50 and P65, no significant fixed effects were retained in any model. This indicates that these sensor-level TEPs were largely insensitive to pulse type, resting-state or task-related GABA+/Glx measures, or their interactions. In contrast to SM1, where early to mid-latency TEP components (P30, N45, and P60) were robustly predicted by the task-related GABAergic tone, the lack of predictive effects in DLPFC may reflect lower local coupling between neurometabolite levels and TEP generation.

Whereas the GMFP analyses mostly concurred with the sensor-level results for N50, with only a significant effect of resting-state GABA+ (more GABA+ related to a larger, more negative, N50 amplitude), the effects of P65 were considerably different (cf., **Supplementary Materials 3**). Specifically, several neurometabolite-predictors were significant. Resting-state and task-related Glx showed opposite relationships depending on TMS condition: higher Glx was associated with smaller P65 amplitudes in the single-pulse condition, whereas in the paired-pulse (LICI) condition, higher Glx was linked to larger P65 amplitudes. Because these effects were not seen in the primary sensor-level analyses, they may reflect model-specific variability rather than robust physiological relationships. Moreover, interpreting whether these GMFP-specific effects represent genuine physiological relationships is further complicated by the fact that DLPFC components, such as P65, are not canonical components in the same way as those in SM1.

#### 3.2.2. The N100 component

For N100, two significant sensor-level findings emerged. First, across all models, N100 amplitude was larger in the single-pulse compared to the paired-pulse condition (p < 0.001, F_1,14_ = 20.632), echoing a trend also observed in SM1 which only reached significance on the GMFP level (**Supplementary Materials 3 and 5**). Second, model 3 revealed a significant interaction between pulse type and the GABA+/Glx ratio. In the paired-pulse, LICI, condition, higher GABA+/Glx ratios were associated with smaller (less negative) N100 amplitudes, whereas in the single-pulse condition, higher GABA+/Glx ratios predicted larger N100 amplitudes (p = 0.008, F_1,13_ = 9.78). The GMFP analyses mostly concurred with the sensor-level analyses, with a consistent effect of pulse type.

These results suggest that in participants with higher inhibitory tone, the paired-pulse paradigm induced stronger suppression, leading to a smaller (less negative) N100 response, compared to single-pulse TMS. Conversely, in the single-pulse condition, N100 amplitude was largest among participants with higher inhibitory tone.

### 3.3. Intra-measurement correlations

Exploratory correlations among TEP components and MRS neurometabolites are reported in **Supplementary Materials 4**.

For SM1, the observed patterns were consistent with the prior results, where associations between P30, N45 and P60 and GABAergic processes were largely concordant, and N100 emerged as an outlier. Here, we found clustering of early to mid-latency TEP components (P30–P60) with each other but not N100, in addition to internal coherence among MRS metabolites.

For DLPFC, exploratory correlations revealed lower interrelatedness than observed in SM1. Only task-related and resting-state GABA+, and task-related and resting-state Glx correlated significantly, with no cross-metabolite relationships. Consistent with SM1, for TEPs, N100 did not correlate with earlier DLPFC components (N50, P65), whereas N50 and P65 were significantly correlated with each other.

## 4. Discussion

The present work aimed to link local cortical excitability responses to local neurometabolite concentrations. By combining single- and paired-pulse TMS-EEG over SM1 and DLPFC with both task-related and resting-state MRS, we systematically characterized how neurometabolites relate to TEP components across contexts. Overall, the findings demonstrate region-, brain-state and TEP component specific links. As hypothesized, SM1 results revealed a robust relationship between task-related GABA+ and early to mid-latency SM1 TEPs (P30, N45 and P60), independent of the employed TMS paradigm. Conversely, more complex associations were found for DLPFC and the later N100 component (**Figure 6**).

**Figure 6.**
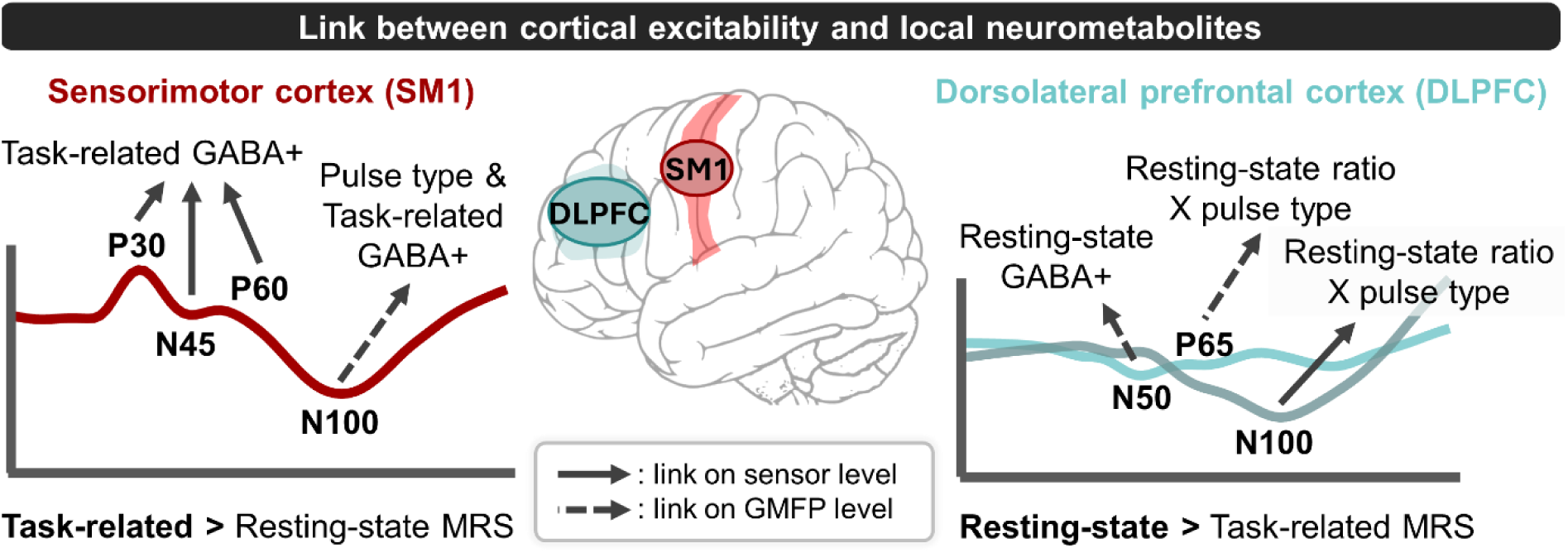
Overview of the main results. Per transcranial evoked potential (TEP) component, if significant sensor level results were present, these are visualized irrespective of global mean field power (GMFP) results as full lines, as the local link between neurometabolites and cortical excitability responses was our primary focus. If these local results were not significant and significant GMFP results were present, these are presented with dashed lines. Likewise, for each component, only the significant effects from the model that explained the most variance is shown. This does not imply that other neurometabolite measures did not explain TEPs, as, for instance, SM1 P30 was also explained by resting-state Glx, albeit in a lesser manner than task-related GABA+.

### 4.1. SM1: Task-related GABAergic tone shapes local TEPs

In SM1, early to mid-latency TEP components (P30, N45, and P60) were consistently best predicted by task-related GABA+ levels, with higher inhibitory tone relating to larger TEP amplitudes across all three components. This aligns with our hypotheses and builds on prior pharmacological work demonstrating the sensitivity of several TEPs components to GABA**-**mediated inhibition (Premoli *et al*., 2014a; Darmani & Ziemann, 2019). Importantly, task-related MRS outperformed resting-state MRS in predicting these components, suggesting that inhibitory dynamics during task-engagement are a more predictive read-out of cortical reactivity than baseline neurometabolite levels alone.

The finding that GABA+ outperformed the GABA+/Glx ratio for these early to mid-latency components further indicates that local GABAergic inhibitory tone exerts a dominant influence on SM1 reactivity in the first ∼60 ms following stimulation. In accordance with prior work, a strong relationship between neurometabolites and the SM1 N100 was absent (Du *et al*., 2018). Together, these results underscore a hierarchy of neurometabolite-sensitivity in TEP components. While earlier, more local, TEP components reflect inhibitory processes of reverberant activity over SM1, the later N100 component may reflect a broader, multisensory, network-level response (Farzan & Bortoletto, 2022). This interpretation is further supported by the presence of a significant TEP-neurometabolite relationship for N100 in the global field, but not the local sensor, analyses.

Overall, these SM1 findings extend beyond prior electromyography-based TMS research by establishing several canonical local TEP components as accessible indices of task-related GABAergic inhibitory tone in SM1. Going forward, these components may hold value as biomarkers for (i) conditions characterized by aberrant GABA concentrations, (ii) the personalization of neuromodulation interventions, and (iii) the evaluation of treatment efficacy.

### 4.2. DLPFC: Complex sensitivity to local neurometabolites

In contrast to SM1, DLPFC TEPs were largely insensitive to both resting-state and task-related MRS measures. Early components (N50, P65) showed no significant associations in the local sensor analyses, while N100 revealed a pulse type × resting-state GABA+/Glx interaction and a main effect of pulse type. Single-pulse TMS over DLPFC revealed that a higher N100 amplitude was related to a larger amount of GABA+ relative to Glx, an opposite relationship to that identified in prior work (Du *et al*., 2018). Moreover, paired-pulse TMS data supported the single-pulse results by showing that participants with a strong inhibitory tone had a greater inhibited response (i.e., more LICI), quantified as a lower (less negative) amplitude of the N100 component.

Several factors may explain the regional discrepancy between SM1 and DLPFC in terms of linking TEPs to neurometabolites. For one, sensor-level and GMFP results converged in showing that, in the DLFPC, resting-state neurometabolite measures provided a better fit, whereas in SM1, models containing task-related measures consistently showed the better fit. While the employed motor coordination task, the BTT, recruits prefrontal areas (Swinnen & Wenderoth, 2004; Beets *et al*., 2015; Santos Monteiro *et al*., 2017; Zivari Adab *et al*., 2020), it remains primarily a motor task, engaging SM1. Task-related MRS effects in DLPFC may therefore have been attenuated due to limited recruitment of inhibitory-excitatory circuits. If so, this suggests that stronger associations between DLPFC excitability and neurometabolites may emerge when using a task more directly engaging DLPFC, such as a working memory paradigm (Ragland *et al*., 2002; Razza *et al*., 2024).

In addition, although DLPFC targeting was individualized via fMRI, we aligned the stimulation dose and coil orientation with the parameters used for SM1 to remain consistent with established best-practice guidelines. Nevertheless, this does not consider genuine inter-regional differences in anatomy and neurophysiology (Kolk & Rakic, 2022; Van Hoornweder *et al*., 2024). Real-time TMS-EEG optimization for DLPFC may help to account for some of these factors, by optimizing the stimulation dose and coil orientation to local TEP components (Casarotto *et al*., 2022; Gogulski *et al*., 2024). Likewise, precise adjustments in coil positioning may reduce TMS-induced blinks and decay artifacts, thereby decreasing the number of excluded DLPFC datasets. In our study, the smaller number of participants with high-quality DLPFC recordings likely limited statistical power to detect subtle effects.

### 4.3. Pulse type: Paired-pulse offers limited sensitivity beyond single-pulse measures

Across both regions, we applied single-pulse TMS and a paired-pulse, LICI, paradigm, probing GABA_B_ receptor activity (Valls-Solé *et al*., 1992; Werhahn *et al*., 1999; Premoli *et al*., 2014b). We did not observe a main effect of pulse type on local TEP components for both SM1 (P30, N45, and P60) and DLPFC (N50, and P65) (cf. **Supplementary Materials 5**). Conversely, the N100 component was consistently affected by pulse type. Overall, this concurs with prior pharmacological studies over SM1 which link earlier TEP components to GABA_A_, and later components to GABA_B_ activity (Premoli *et al*., 2014a; Darmani & Ziemann, 2019). However, the hypothesis that pulse type-related modulation of the N100 depends on GABA+, assuming both reflect GABA_B_-mediated processes, was only supported for DLPFC.

The current results suggest that prior reports of TEP suppression in LICI paradigms, assessed across the whole brain or entire TEP time-window (Farzan *et al*., 2010; Poorganji *et al*., 2023; Takano *et al*., 2025), may have been driven by a significant global suppression of the late N100 component. In accordance with our results, (de Goede *et al*., 2020) found robust N100 suppression at multiple LICI interstimulus intervals, but no suppression for any local early to mid-latency SM1 TEPs.

Interpretation of the N100 component should be done with caution. Unlike early to mid-latency TEPs, N100 mostly lacked sensor-level correlations with local MRS metabolites or other TEPs, but did display a pulse type effect, particularly at the GMFP level. This implies that N100 reflects distributed, network-level processing that integrates recurrent cortical activity and multisensory afference (Conde *et al*., 2019; Farzan & Bortoletto, 2022). GMFP, by summing absolute field strength across the scalp, enhances sensitivity to spatially distributed sources and therefore revealed paired-pulse modulation that single sensor analyses mostly missed. Consequentially, the effect of the paired-pulse LICI protocol on N100 may reflect activity related to the processing of ongoing sensory co-stimulation from the conditioning stimulus, by stimulating on top of an ongoing N100 component. Although we follow the current best practice approach that subtracts ongoing single-pulse activity relative to the paired-pulse, there is no method available to disentangle the interaction effect by the superimposition of the test stimulus on the ongoing N100 component of the conditioning pulse (Premoli *et al*., 2014b).

Overall, the present study showed that single-pulse TEPs provide a dynamic index of the inhibitory tone of a stimulated region, specifically SM1, with little additional insights from paired-pulse TMS-EEG measures for local responses. Researchers examining region-specific excitability may therefore wish to carefully weigh the additional methodological and acquisition demands of paired-pulse protocols. When acquisition time is limited, increasing the number of single-pulse trials may yield a more meaningful gain in signal-to-noise ratio for local TEP components than the incremental information offered by paired-pulse paradigms.

### 4.4. Limitations and future recommendations

Beyond the already mentioned considerations and recommendations, such as the DLPFC sample size and real-time optimization approaches, several other factors should be acknowledged. EEG hardware limitations did not enable the recording of the immediate cortical response within the first milliseconds following a TMS pulse (Stango *et al*., 2025). While immediate TEPs and early to mid-latency TEPs follow a similar rostro-caudal response following stimulation over SM1 (Nuyts *et al*., 2025a), studying the immediate responses could establish a more direct link between the stimulated neural circuit and local neurometabolites (Beck *et al*., 2024). In similar vein, studying how local neurometabolite levels relate to the early N15 TEP component, which we were unable to reliably recover in the current dataset, would be interesting. Lastly, a promising avenue would be the application of TMS-EEG paradigms during task-related brain states (Beck *et al*., 2025; Thong *et al*., 2025), in line with our MRS data demonstrating the relevance of brain states for inhibition/excitation dynamics.

## 5. Conclusion

In summary, we found neurometabolite-cortical excitability links to be region and brain-state dependent. In SM1, local early to mid-latency TEPs were robustly related to the region’s task-related GABAergic inhibitory tone, whereas the later, network-level, N100 component showed limited relationships with neurometabolites. DLPFC TMS-derived responses revealed fewer links to local neurometabolites, with most associations relating to (i) resting-state neurometabolite-levels and (ii) the global field TEP level. Pulse type (single-vs paired-pulse TMS) almost exclusively impacted the N100 component, suggesting that single-pulse TEPs may already capture a large extent of the relevant variation in the studied neurometabolites at the cortical level. More broadly, these results provide some of the most compelling evidence to date linking MRS-derived neurometabolites with TMS-EEG markers of cortical excitability. In doing so, we position several TEP components as accessible, non-invasive markers of inhibitory and excitatory processes in the human cortex. This has implications for basic neuroscience, biomarker development, and the design of individualized neuromodulation interventions aimed at restoring or enhancing cortical function.

## CrediT Author Contribution Statement

Conceptualization: MN, SC, SPS, SVH. Methodology: SC, SPS. Software: MN, SC, SVH. Validation: MN, SC, GRN, SVH. Formal analysis: MN, SVH. Investigation: SC. Resources: SC, SPS. Data curation: MN, SC, GRN, SVH. Writing – Original Draft: MN, SVH. Writing – Review & Editing: MN, SC, GRN, KC, RM, SPS, SVH. Visualization: MN, SVH. Supervision: SPS, SVH. Project administration: SC, SPS. Funding acquisition: SC, SPS.

