## Supplementary Materials for "Resting-state and task-related neurometabolite levels differentially relate to cortical excitability in the sensorimotor and prefrontal cortex"

[*] Shared first authorship

[†] Shared last authorship

Address for correspondence:

Marten Nuyts

Hasselt University, Faculty of Rehabilitation Sciences

Universiteit Hasselt, Campus Diepenbeek, Faculteit Revalidatiewetenschappen, Wetenschapspark 7, B-3590 Diepenbeek

ORCID-ID: 0000-0002-7687-9897

&

Sybren Van Hoornweder

Hasselt University, Faculty of Rehabilitation Sciences

Universiteit Hasselt, Campus Diepenbeek, Faculteit Revalidatiewetenschappen, Wetenschapspark 7, B-3590 Diepenbeek

ORCID-ID: 0000-0002-0325-8950

**Supplementary materials 1. Task setups.**

**N-back task**

The N-back task is a widely used working memory task to probe activation in specific brain regions, including the dorsolateral prefrontal cortex (DLPFC) (Ragland *et al.*, 2002). During this task, participants are presented with a series of letters and must recall whether the current letter on the screen matches the one 'N' steps earlier. In the present study, participants completed three blocks of the 0-back task, responding whenever the letter 'X' appeared, and three blocks of the 2-back task, in which they responded when the current stimulus matched the one presented two steps back in the series (cf., **Figure SM1.1**). Participants pressed a button on a keyboard whenever a target letter was shown. Blocks were pseudorandomized and consisted of 24 letters each, including five target letters. Before each block, a brief instruction was shown (2 s). Each letter was presented for 1 s with an interval of 1 s.

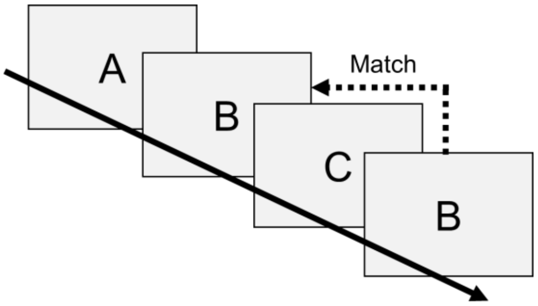


**Figure SM1.1**. N-back task. The paradigm presents a 2-back version where a target match is presented from 2 steps back in the series.

**Bimanual coordination task (BTT)**

The bimanual tracking task (BTT) assesses bimanual motor coordination and was used here to examine task-related changes in neurometabolite levels with magnetic resonance spectroscopy (MRS) (Sisti *et al.*, 2011).

While lying supine in the scanner, participants viewed the task via a mirrored projection screen positioned at eye level (LCD projector: Barco 6300, 1280 × 1024 pixels). A non-ferromagnetic BTT setup was placed over their thighs. During the task, participants tracked a moving dot along a target line by coordinatively moving both hands, each controlling one axis via a dial (**Figure SM1.2**). The right hand controlled movement along the X-axis and the left hand along the Y-axis. Each trial started with a blank screen (2 s), followed by a preparation phase (2 s) during which participants were instructed to plan but not initiate movement. This was followed by an execution phase (8 s) in which they tracked a target dot moving outward from the center of the screen (**Figure SM1.2A**). Target lines appeared in one of four quadrants, requiring the fingers to rotate left, right, inward, or outward. Each BTT block consisted of 48 trials.

Importantly, within the broader research protocol, participants completed both a low and high complexity version of the BTT on two separate days in a pseudorandomized order. In the low complexity version (**Figure SM1.2B**), all trials had an interhand frequency ratio of 1:1, meaning both dials were required to rotate at an equal speed. In the high complexity version (**Figure SM1.2C**), trials used interhand frequency ratios of 2:3 or 1:2, requiring the right hand to move faster than the left to match the target trajectory.

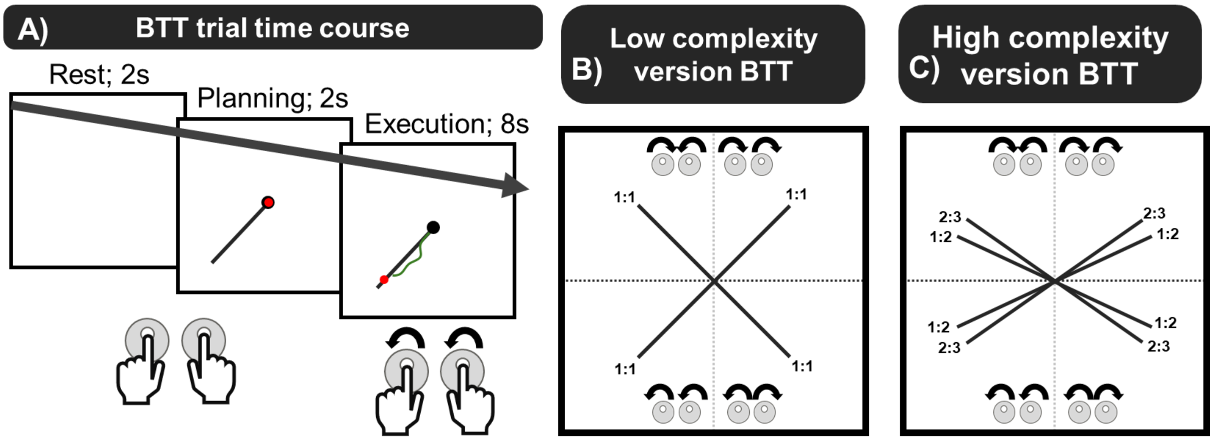


**Figure SM1.2**. The bimanual tracking (BTT). **A)** Exemplary time course of a BTT trial. **B)** Low complexity version of the BTT. **C)** High complexity version of the BTT.

Notably, MRS measures from the high and low complexity versions of the BTT were highly correlated (**Figure SM1.3**). Therefore, we used the data collected during the high complexity version, as the session involving the low complexity version had a higher participant dropout (n = 2).

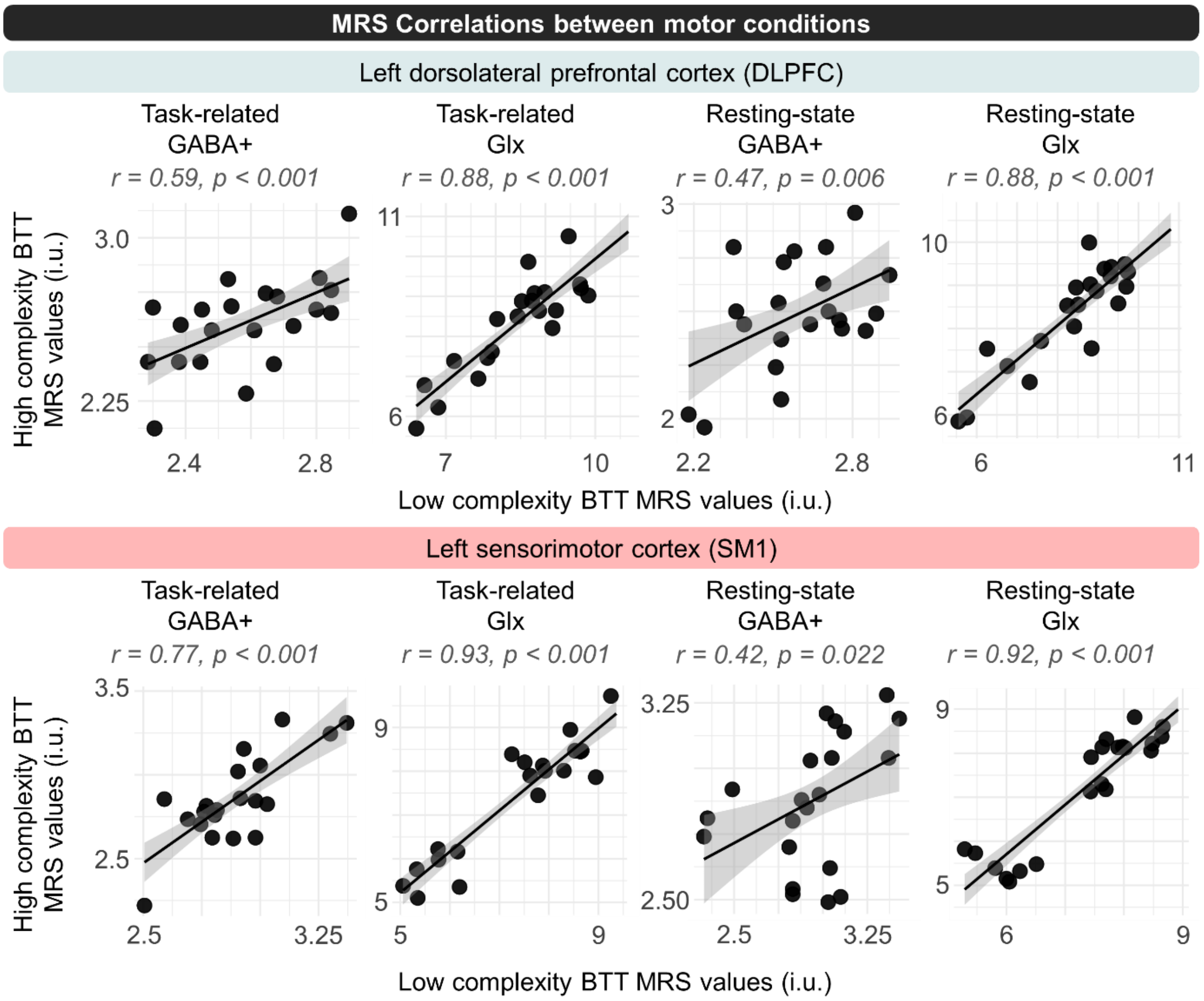


**Figure SM1.3.** Correlations between neurometabolite estimates across the two bimanual tracking task (BTT) sessions. In the main manuscript, we used the high complexity BTT data (y-axis) because this condition yielded the largest number of valid datasets. Across both regions—dorsolateral prefrontal cortex (DLPFC) and primary sensorimotor cortex (SM1)—all neurometabolites showed significant between-session correlations. This consistency indicates that the primary findings would likely remain unchanged had the low complexity BTT condition been used instead. Correlation p-values were Bonferroni-corrected for four comparisons per region.

### Supplementary Materials 2. MRS reporting

This section provides an overview of the minimal reporting standards for the in vivo MRS, as described in (Lin *et al.*, 2021).

| Site (Name or Number) | |
|---|---|
| 1. Hardware | |
| a. Field strength [T] | 3 Tesla |
| b. Manufacturer | Philips |
| c. Model (software version if available) | Achieva |
| d. RF coils: nuclei (transmit/ receive), number of channels, type, body part | 32-channel receive-only head coil (Philips), body transmit; nucleus: 1H |
| e. Additional hardware | / |
| 2. Acquisition | |
| a. Pulse sequence | GABA-edited MEGAPRESS sequence |
| b. Volume of Interest (VOI) locations | Left DLPFC and left SM1 |
| c. Nominal VOI size [cm3, mm3] | Left DLPFC - voxel size =, 40 × 25 × 25 mm3<br>Left SM1 - voxel size = 30 × 30 × 30 mm3 |
| d. Repetition Time (TR), Echo Time (TE) [ms, s] | TR = 2000 ms, TE = 68 ms |
| e. Total number of Excitations or acquisitions per spectrum<br>In time series for kinetic studies<br>i. Number of Averaged spectra (NA) per time-point | DLPFC: 160 averages (~ 11 min)<br>SM1: 112 average (~ 8 min) |

| | |
|---|---|
| ii. Averaging method (e.g. block-wise or moving average)<br>iii. Total number of spectra (acquired / in time-series) | |
| f. Additional sequence parameters<br><br>(spectral width in Hz, number of spectral points, frequency offsets)<br><br>If STEAM:, Mixing Time (TM)<br><br>If MRSI: 2D or 3D, FOV in all directions, matrix size, acceleration factors, sampling method | Flip angle = 90°, 2 kHz spectral width, 'ON'/'OFF' editing pulse frequency = 1.9 / 7.5 ppm, 'ON'/'OFF' spectra collected in an interleaved fashion |
| g. Water Suppression Method | "MOIST" |
| h. Shimming Method, reference peak, and thresholds for "acceptance of shim" chosen | Automatic shimming was performed before every MRS acquisition, and 16 unsuppressed water averages were obtained over each brain region of interest and later used for referencing. |
| i. Triggering or motion correction method<br><br>(respiratory, peripheral, cardiac triggering, incl. device used and delays) | None |
| **3. Data analysis methods and outputs** | |
| a. Analysis software | Gannet (version 3.2.1) |
| b. Processing steps deviating from quoted reference or product | • Frequency and phase correction using spectral registration.<br>• Fitting the difference spectrum (ON–OFF) with a three-Gaussian model between 2.8–4.2 ppm.<br>• Water modeling with Lorentz-Gaussian lineshape. |

| | |
|---|---|
| | • Coregistration and segmentation for partial-volume correction (GannetCoRegister, GannetSegment).<br>• Using high-resolution T1 from Day 2 co-registered to the short anatomical for improved segmentation accuracy.<br>• Tissue-fraction correction via Equation 5 (Harris et al., 2015) using GannetQuantify.<br>• Visual QC procedures (water suppression, lipids, fit error, SNR). |
| c. Output measure<br><br>(e.g. absolute concentration, institutional units, ratio) | Institutional units |
| d. Quantification references and assumptions, fitting model assumptions | Water signals modeled with a Lorentz–Gaussian lineshape were used as the internal reference for metabolite quantification. Difference-edited spectra were fitted using a three-Gaussian model for the GABA+/Glx region (2.8–4.2 ppm), constituting the assumed lineshape and peak composition underlying concentration estimates. Tissue-fraction corrections were applied following Equation 5 of Harris et al. (2015), using grey-matter, white-matter, and cerebrospinal-fluid fractions obtained through segmentation. The high-resolution Day-2 T1-weighted anatomical scan was used as the structural reference for segmentation, assuming improved tissue classification accuracy relative to the short anatomical scans used during voxel placement. The reported GABA+ values are macromolecule-containing. |
| 4. Data Quality | |
| a. Reported variables<br><br>(SNR, Linewidth (with reference peaks)) | fit error, signal-to-noise ratio, Linewidth |
| b. Data exclusion criteria | fit error > 10; high water frequency fluctuations; |
| c. Quality measures of postprocessing Model fitting (e.g. CRLB, goodness of fit, SD of residual) | Goodness of fit |
| d. Sample Spectrum | Included in figure 2 (SM1) and figure 4 (DLPFC) of the main manuscript. |

**Supplementary Materials 3. Global Mean Field Power (GMFP) Results**

To complement the sensor-level peak analyses, we repeated all statistical models using global mean field power (GMFP) data of which the results are reported below.

**Table SM3.1.** Results of models examining the link between SM1 TEP (GMFP) components and neurometabolites.

| TEP | Model | Predictors Retained | AIC | BIC | $R^2_{marginal}$ | $R^2_{conditional}$ |
|---|---|---|---|---|---|---|
| **P30** | **2** | **Task-related GABA+ * Pulse Type**<br>***p* = 0.037, *F* = 5.05**<br>**Task-related GABA+**<br>***p* = 0.007, *F* = 9.03**<br>***Pulse Type***<br>***p* = 0.043, *F* = 4.78** | **91.75** | **101.88** | **0.310** | 0.901 |
| | 1, 3, 4 | / | 99.12 | 104.19 | 0 | 0. 880 |
| N45 | 1, 3 | None | 113.42 | 118.49 | 0 | 0.887 |
| | **2** | **Task-related GABA+**<br>***p* = 0.002, *F* = 12.65** | **104.78** | **111.53** | **0.372** | 0.889 |
| | 4 | Task-related ratio<br>*p* = 0.026, *F* = 5.88 | 109.77 | 116.52 | 0.219 | 0.890 |
| P60 | 1, 3 | None | 120.41 | 125.47 | 0 | 0.819 |
| | **2** | **Task-related GABA+**<br>***p* = 0.002, *F* = 12.95** | **111.57** | **118.32** | **0.364** | 0.823 |
| | 4 | Task-related ratio<br>*p* = 0.042, *F* = 4.81 | 117.67 | 124.42 | 0.181 | 0.826 |
| N100 | 1, 3, 4 | Pulse Type<br>*p* < 0.001, *F* = 24.12 | 121.91 | 128.66 | 0.218 | 0.647 |
| | **2** | **Pulse Type**<br>***p* < 0.001, *F* = 24.12**<br>**Task-related GABA+**<br>***p* = 0.009, *F* = 8.47** | **116.19** | **124.64** | **0.400** | **0.653** |

**The significant interaction between task-related GABA+ and Pulse Type for P30 in model 2 entailed a slightly stronger relationship between GABA+ and P30 amplitude (GMFP) for Single-Pulse TMS than Paired-Pulse TMS.*

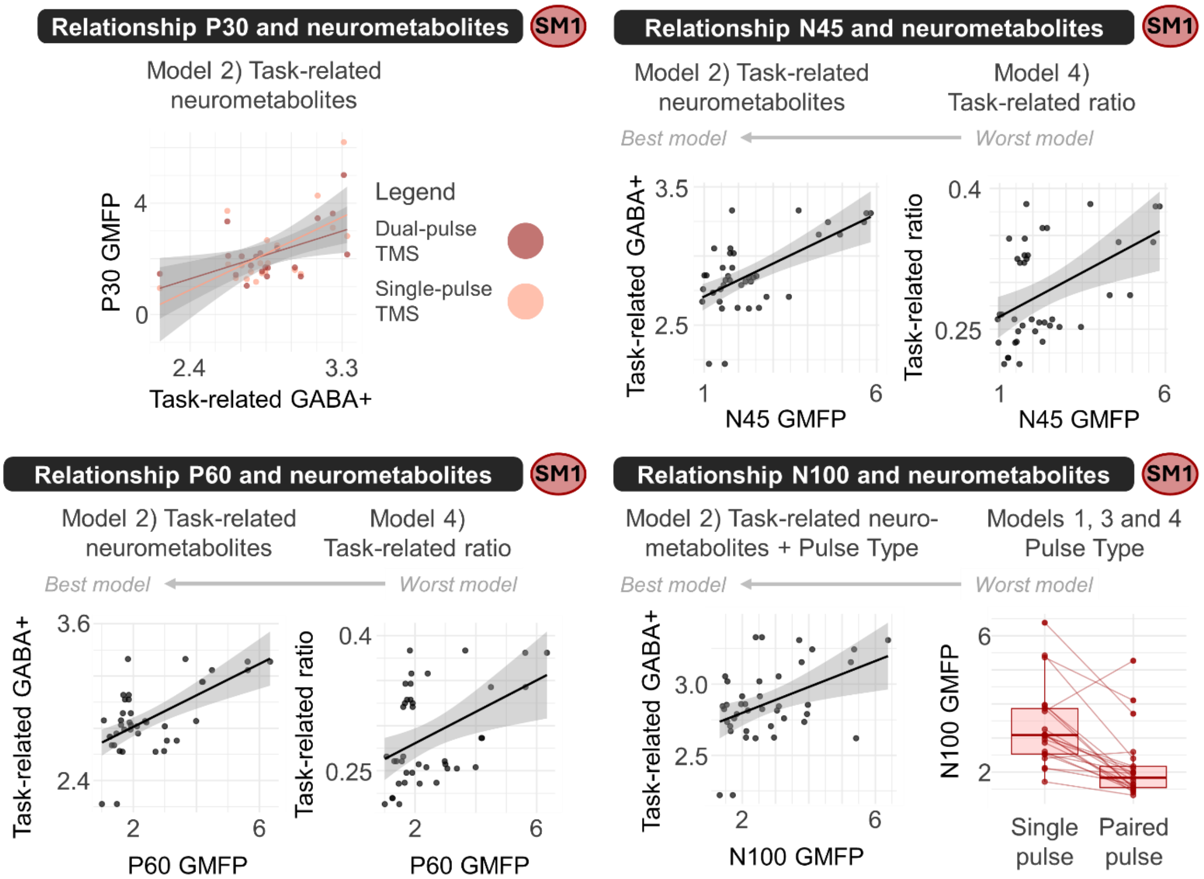


**Figure SM3.1.** Main results of sensorimotor cortex (SM1) global mean field power (GMFP) analyses (cf., Table SM3.1.).

**Table SM3.2**. Results of models examining the link between DLPFC TEP (GMFP) components and neurometabolites.

| TEP | Model | Predictors Retained | AIC | BIC | $R^2_{marginal}$ | $R^2_{conditional}$ |
|---|---|---|---|---|---|---|
| N50 | 2 - 4 | None | 55.28 | 59.48 | 0 | 0.487 |
| | **1** | ***Resting-state GABA+***<br>***p = 0.015, F = 7.91*** | **50.15** | **55.75** | **0.265** | **0.500** |
| **P65** | **1** | **Resting-state Glx*Pulse Type**<br>**p = 0.021, F = 6.88**<br>**Resting-state Glx**<br>**p = 0.219, F = 9.85**<br>**Pulse Type**<br>**p = 0.011, F = 8.80**<br>**Resting-state GABA+**<br>**p = 0.009, F = 9.85** | **51.89** | **61.69** | **0.434** | **0.598** |
| | 2 | Task-related Glx*Pulse Type<br>p = 0.016, F = 7.65<br>Task-related Glx<br>p = 0.563, F = 0.35<br>Pulse Type<br>p = 0.008, F = 9.676 | 57.91 | 66.31 | 0.208 | 0.613 |
| | 3 | None | 63.54 | 67.74 | 0 | 0.233 |
| | 4 | Task-related ratio*Pulse Type<br>p = 0.024, F = 6.48<br>Task-related ratio<br>p = 0.806, F = 0.81<br>Pulse Type<br>p = 0.051, F = 4.63 | 59.11 | 67.51 | 0.185 | 0.591 |
| N100 | **1 - 4** | **Pulse Type**<br>**p > 0.001, F = 20.632** | **122.15** | **127.75** | **0.330** | **0.537** |

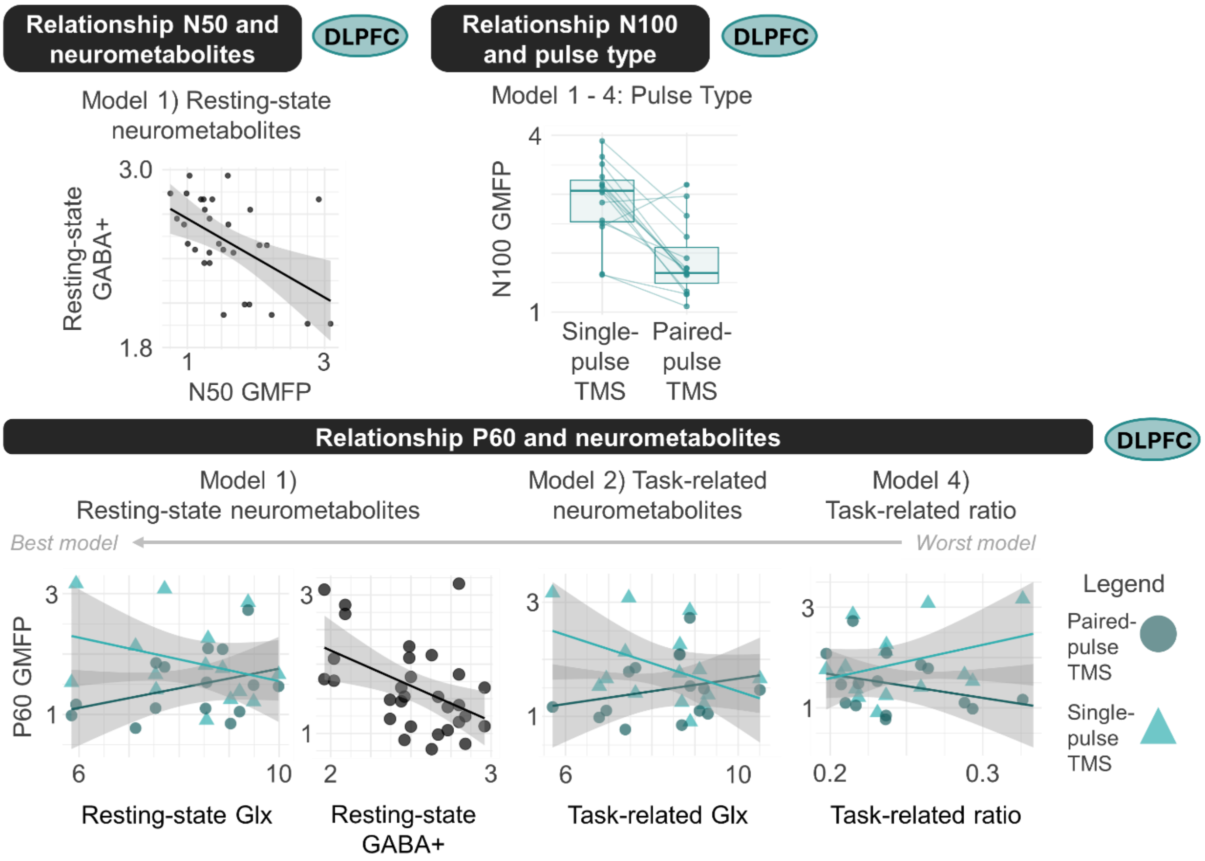


**Figure S2.2.** Main results of dorsolateral prefrontal cortex (DLPFC) global mean field power (GMFP) analyses (cf., Table SM3.2.).

Supplementary Materials 4. Intra-measurement correlations.

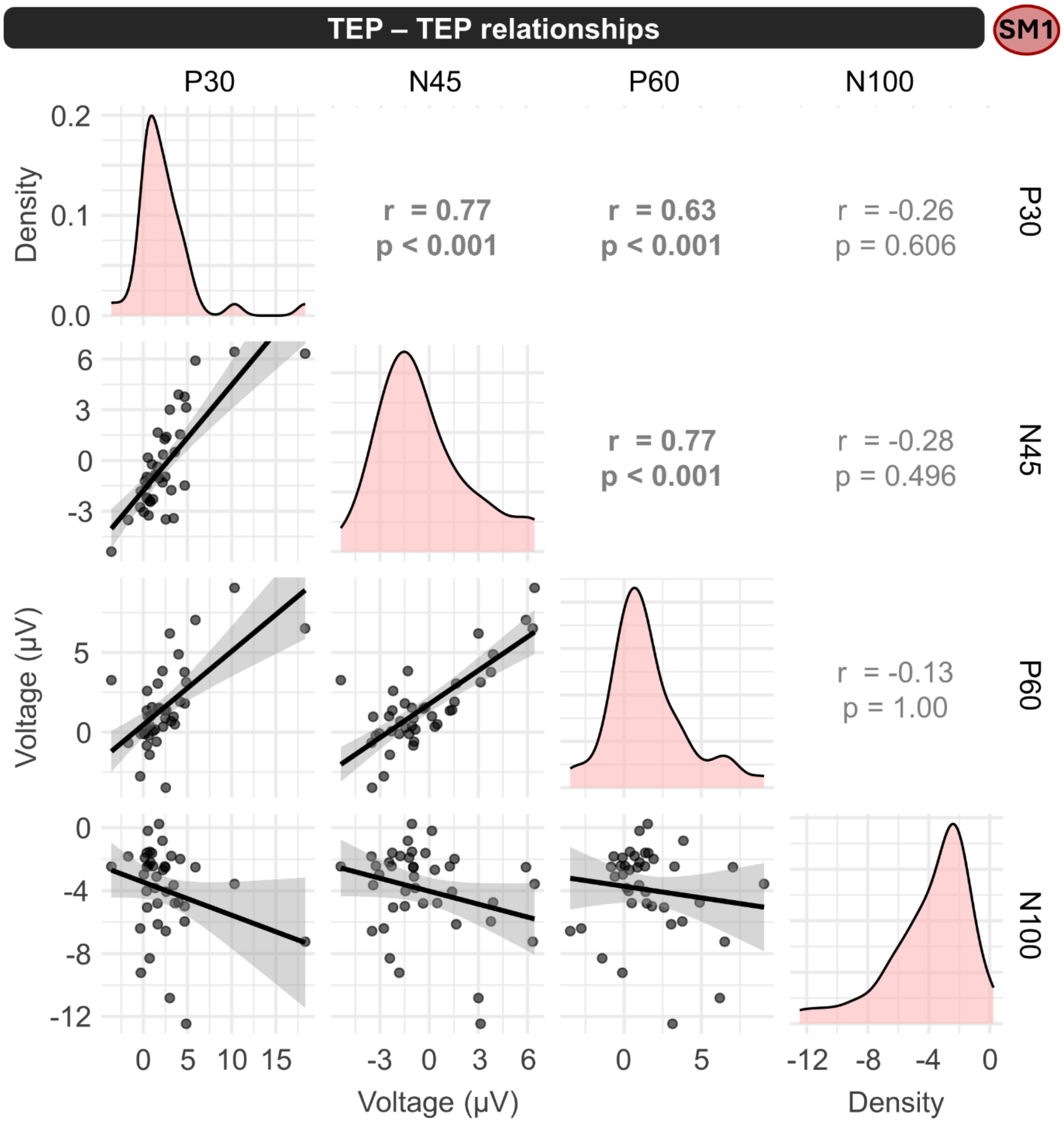


**Figure SM4.1.** Transcranial evoked potential (TEP) - TEP correlations in primary sensorimotor cortex (SM1), alongside the data distribution of each TEP. Pearson correlations were calculated for all pairwise combinations of TEP components and Bonferroni-corrected for six comparisons. Significant correlations were observed between P30, N45, and P60, which also showed similar relationships with SM1 neurometabolites. In contrast, N100 did not significantly correlate with the other components. Importantly, while these results suggest that P30, N45, and P60 are related, within-subject factors such as noise and sensory co-stimulation are also correlated across components and likely inflate correlation coefficients.

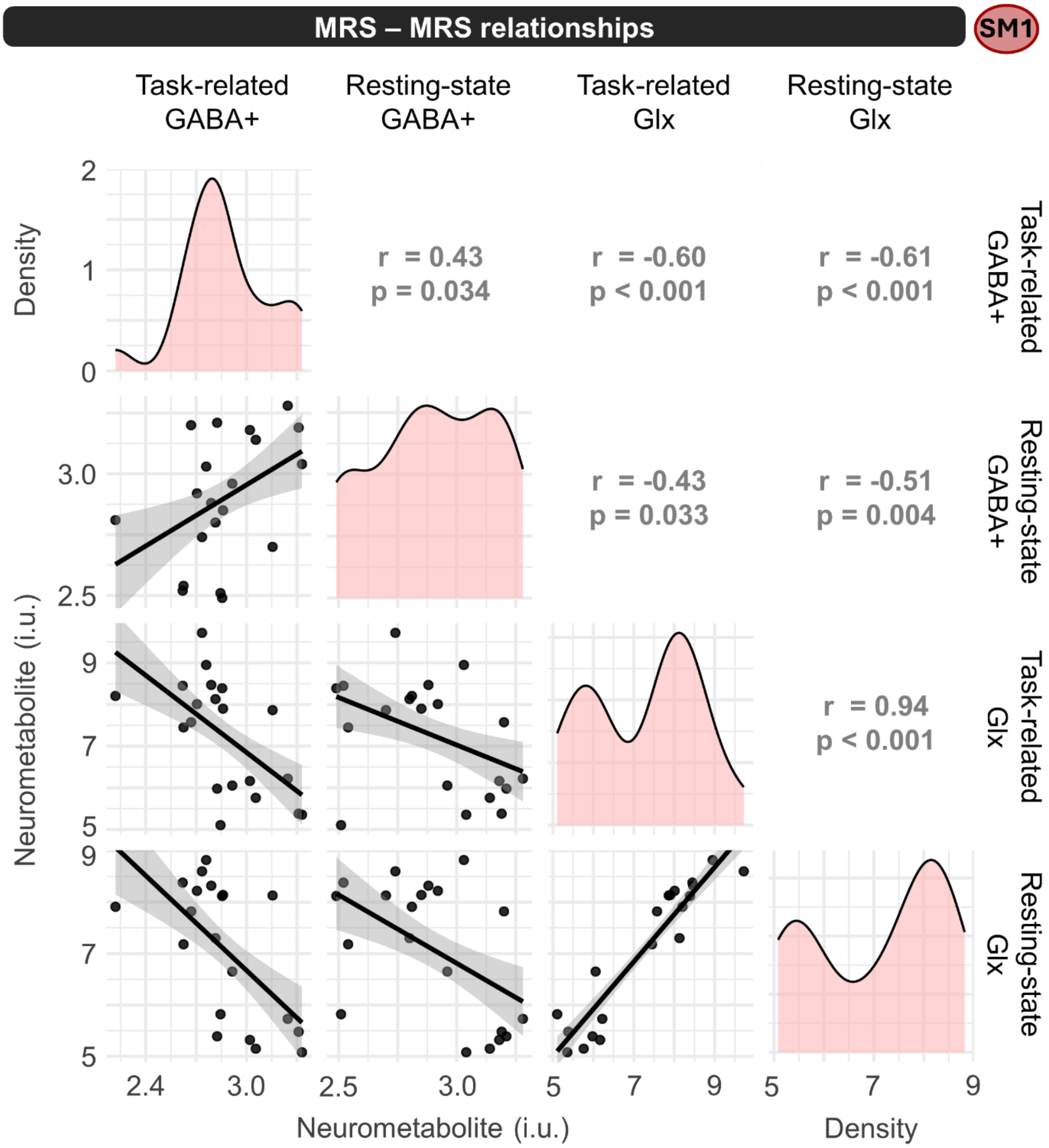


**Figure SM4.2.** Neurometabolite - neurometabolite correlations in primary sensorimotor cortex (SM1), alongside the data distribution of each neurometabolite. Pearson correlations were calculated for all pairwise combinations and Bonferroni-corrected for six comparisons. Significant correlations were observed between all neurometabolites.

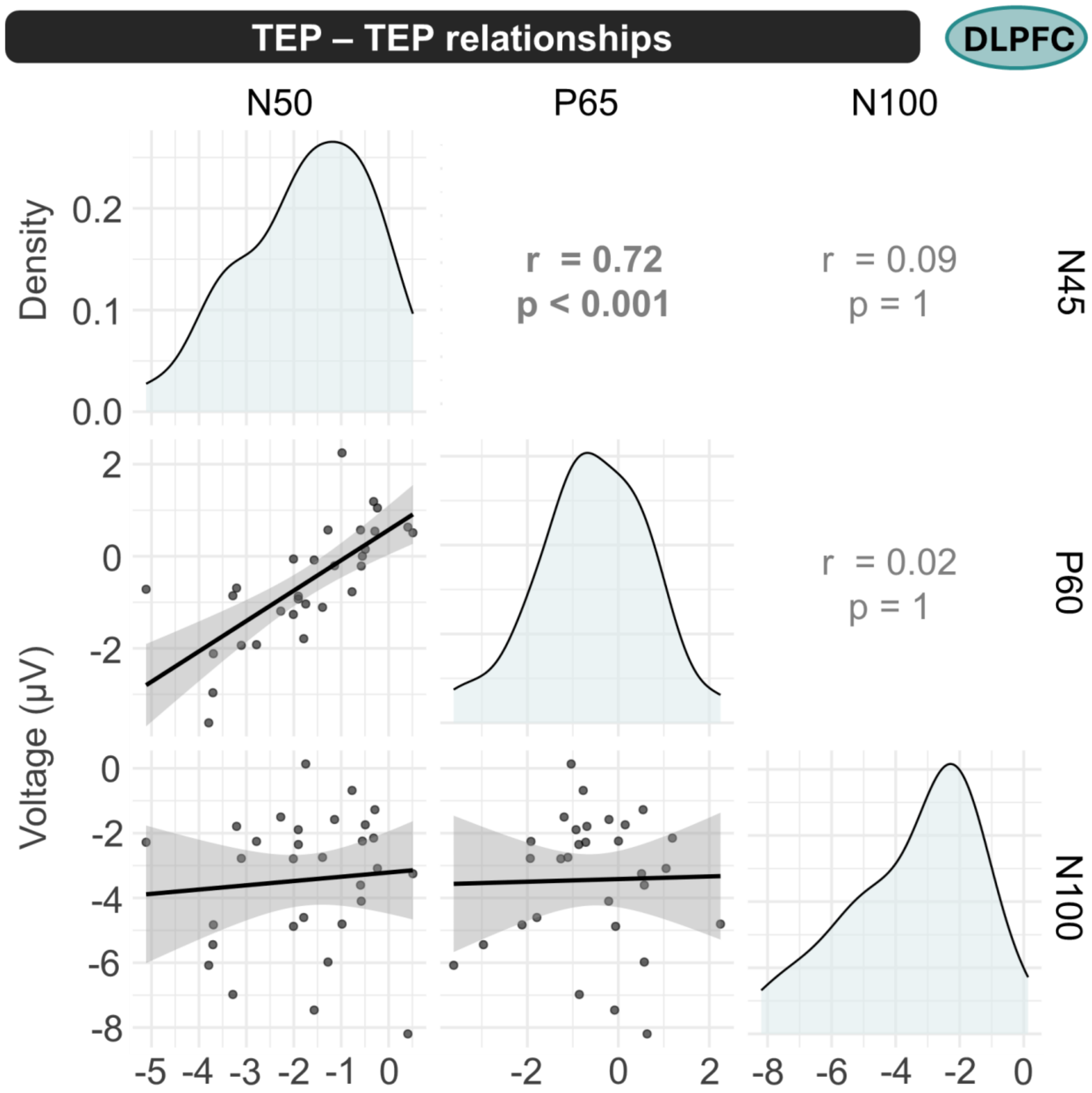


**Figure SM4.3.** Transcranial evoked potential (TEP) - TEP correlations in the dorsolateral prefrontal cortex (DLPFC), alongside the data distribution of each TEP. Pearson correlations were calculated for all pairwise combinations of TEP components and Bonferroni-corrected for three comparisons. Significant correlations were observed between N50 and P60, which also showed a similar absence of association with DLPFC neurometabolites. In contrast, N100 did not significantly correlate with the other components and did show significant effects of neurometabolites and TMS pulse type.

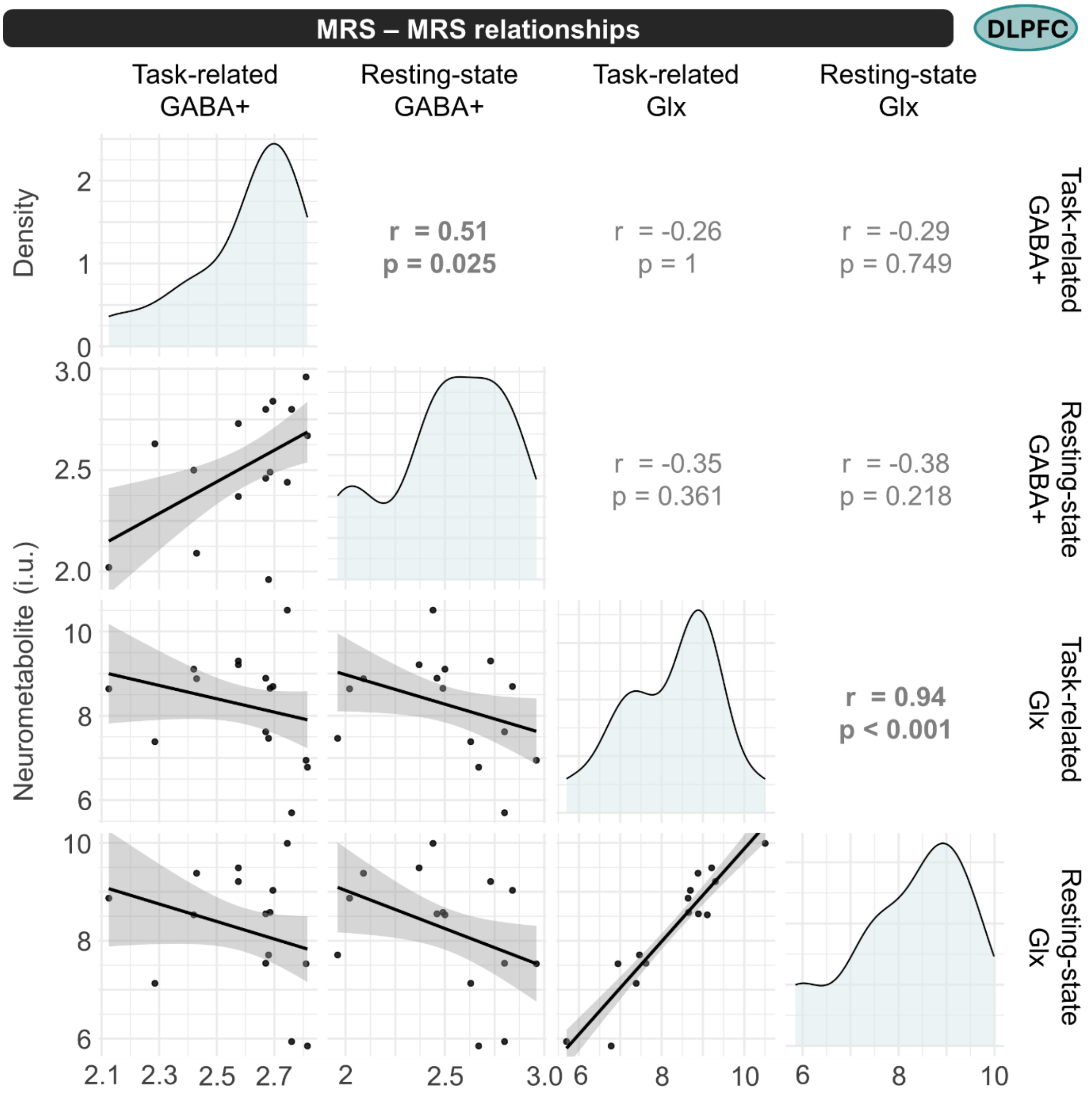


**Figure SM4.3.** Neurometabolite - neurometabolite correlations in the dorsolateral prefrontal cortex (DLPFC), alongside the data distribution of each neurometabolite. Pearson correlations were calculated for all pairwise combinations and Bonferroni-corrected for six comparisons. Significant correlations were only within the specific neurometabolites, across the resting-state and task-related magnetic resonance spectroscopy (MRS) measurements but not across neurometabolites.

**Supplementary Materials 5. Effect of pulse type on sensor-level TEP data.**

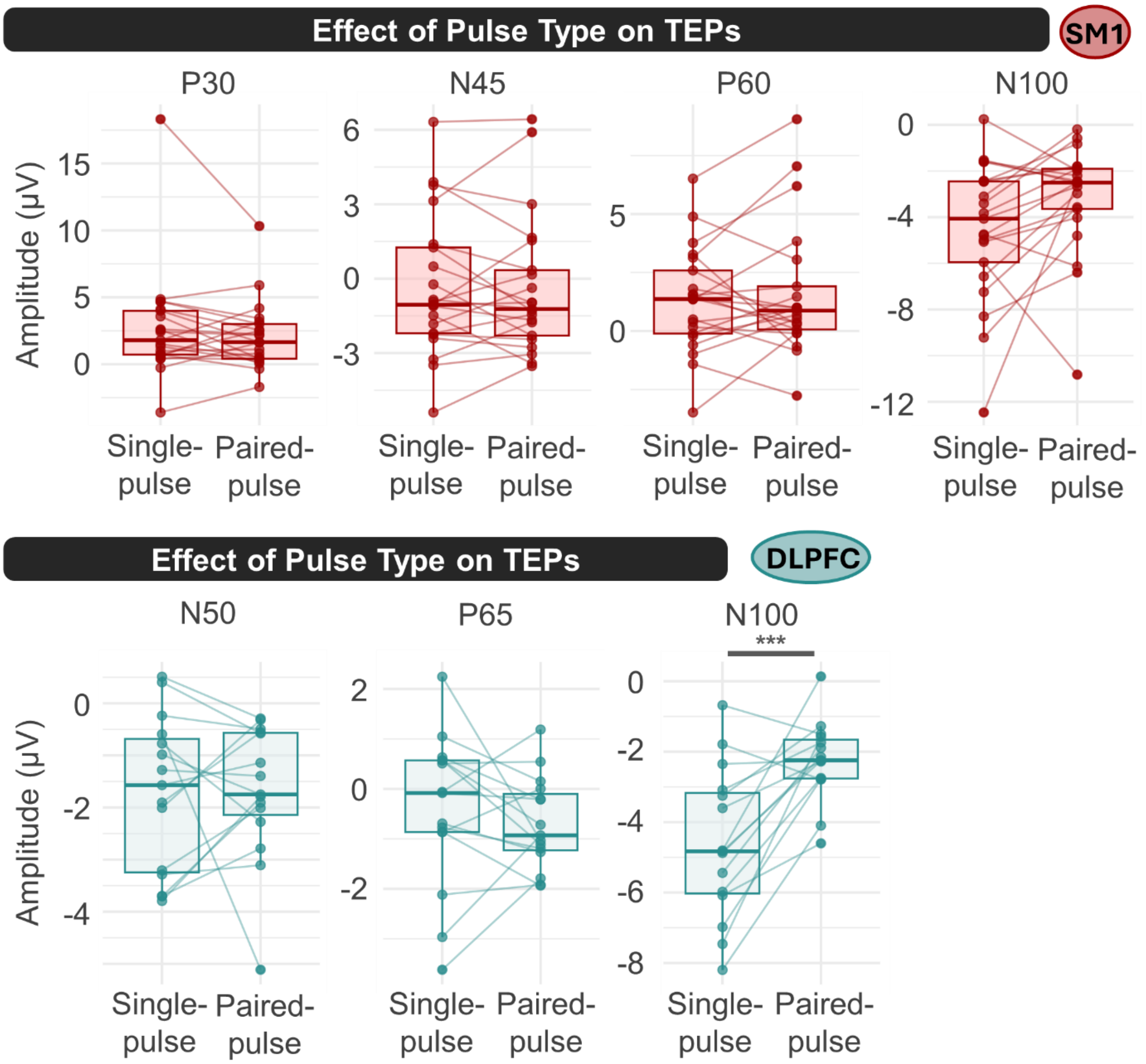


**Figure SM5.1.** Boxplots indicating the difference in transcranial evoked potential (TEP) component amplitudes per component. Only for N100 in dorsolateral prefrontal cortex (DLPFC), a significant effect of pulse type was observed, with a larger N100 during single-pulse transcranial magnetic stimulation (TMS). For N100 over primary sensorimotor cortex (SM1), a similar trend was present, but this did not reach significance for the sensor-level data. However, for the global mean field potential (GMFP) analyses, which present more global phenomena, a significant effect for pulse type on SM1 N100 was observed (cf., **Supplementary Materials 3**).
